# Dynamics of ASC speck formation during skin inflammatory responses *in vivo*

**DOI:** 10.1101/111542

**Authors:** Paola Kuri, Nicole L. Schieber, Thomas Thumberger, Joachim Wittbrodt, Yannick Schwab, Maria Leptin

## Abstract

Activated danger or pathogen sensors trigger assembly of the inflammasome adaptor ASC into specks, large signalling platforms considered hallmarks of inflammasome activation. Because a lack of *in vivo* tools has prevented the study of endogenous ASC dynamics, we generated a live ASC reporter through CRISPR/Cas9 tagging of the endogenous gene in zebrafish. We see strong ASC expression in the skin and other epithelia that act as barriers to insult. A toxic stimulus triggered speck formation and rapid pyroptosis in keratinocytes *in vivo*. Macrophages engulfed and digested this speck-containing pyroptotic debris. A 3D ultrastructural reconstruction based on CLEM of *in vivo* assembled specks revealed a compact network of highly intercrossed filaments, whereas PYD or CARD alone formed filamentous aggregates. The effector caspase is recruited through PYD, whose overexpression induced pyroptosis, but after substantial delay. Therefore, formation of a single compact speck and rapid cell death induction *in vivo* requires full-length ASC.

**One Sentence Summary:** With a new endogenous ASC real-time reporter we characterize speck dynamics *in vivo* as well as the concomitant pyroptosis speck formation causes in keratinocytes.

## Introduction

Inflammasomes are large supramolecular structures that signal the detection of danger or pathogenic stimuli by pattern recognition receptors, including some NOD-like receptor (NLR) family members (1, 2). Inflammasome signalling ultimately leads to the activation of the effector caspase-1 through proximity-induced, auto-proteolytic cleavage (3). Activated caspase-1 can proteolytically process cytokines as well as trigger pyroptosis, a pro-inflammatory form of regulated cell death (4). During pyroptosis, cells swell after pores assemble in the plasma membrane, leading to its rupture and the release of intracellular contents and membrane vesicles (5, 6). The adaptor molecule apoptosis-associated speck-like protein containing a CARD (ASC) is central to the inflammasome assembly process (7). ASC is composed of two protein–protein interaction domains of the death domain superfamily, a pyrin domain and a caspase activation and recruitment domain (PYD_A_ and CARD_A_, respectively) joined by a flexible linker (8). This enables ASC to interact with both PYD-containing receptors and the CARD-containing pro-caspase-1, thus bridging sensor and effector molecules (1).

Upon activation, inflammasome-forming receptors oligomerize and nucleate the prion-like aggregation of ASC, enabling the subsequent clustering of caspase-1(9, 10). During this process, ASC is rapidly depleted from its steady-state homogeneous cellular distribution and self-associates to form a single punctum inside the cell of about 1μm in diameter, called a speck (11, 12). The fast and irreversible assembly of ASC into specks maximizes the amount of activated caspase-1, ensuring a high signal amplification (1, 13).

Structural methods used to analyse specks *in vitro* showed that ASC assembles into filaments of which PYD_A_ forms a rigid cylindrical core while CARD_A_ is directed outwards through a flexible attachment (9, 14). The external orientation of CARD_A_, in addition to enabling the recruitment of downstream signalling elements, allows intra and inter-filament crosslinking through CARD_A_-CARD_A_ interactions. Indeed, recent cell culture studies showed that preventing CARD_A_ interactions by single point mutagenesis (15) or use of an intracellular alpaca antibody fused to a fluorescent protein (16) abolishes speck formation, but not PYD_A_ filament assembly. However, whether *in vivo* assembled specks also share this crosslinked filament arrangement remains to be analysed with structural methods.

By expressing ASC fused to a fluorescent protein from a transgene, specks can be visualized by light microscopy (12, 17). The switch from a diffuse signal throughout the cell to one single bright point is considered a readout and proxy for inflammasome activation (18-21). However, experimentally expressed constructs increase the cellular concentration of ASC and, given the protein’s high tendency to aggregate if overexpressed (7), the risk that speck formation occurs without an inflammatory stimulus also increases. The aforementioned study by Schmidt et al. (2016) represented the first time endogenous ASC was visible in a cell, but because speck formation is abolished by the use of the alpaca antibody, this tool cannot be used to assess speck formation *in vivo*.

Inflammasome function has mainly been studied in cells of the innate immune system such as macrophages. However, many pathogens and toxic agents first enter the body through epithelia that form the interfaces between body and environment, which evidently require innate immune surveillance mechanisms(22), but little is known about the role of the inflammasome and ASC in these, or other tissues such as endothelium or connective tissue which are also composed of cells that contribute to a global inflammatory response (22-24). For example, although ASC is present in mammalian epidermis (25) and it acts as a tumour suppressor in keratinocytes (26), whether speck formation leads to pyroptosis in these cells is unknown. Studying the responses of native tissues *in vivo* using murine models however, is challenging due to limited imaging accessibility. The zebrafish (*Danio rerio*) is a genetically and optically accessible model organism for studying diseases and for drug screening (27-30) in which *in vivo* innate immune responses can be studied in the context of a whole organism (27, 31). The zebrafish genome contains more than ten times as many NLR genes as mice and humans (32-34), but it has only one gene encoding ASC (also called pycard) with a PYD-CARD domain structure (35).

We use zebrafish to study ASC function in tissues, such as the skin, in which inflammasome signalling has not been addressed *in vivo*. The transparency of the zebrafish makes this model especially well suited to study ASC-mediated inflammasome formation using speck formation as readout. For this purpose, we generated a line in which the endogenous *asc* is tagged with GFP using CRISPR/Cas9 technology, allowing body-wide *in vivo* analysis of speck formation.

This tool, together with an *asc* inducible expression system with which we visualize the ultrastructure of specks formed *in vivo*, revealed that speck formation in keratinocytes can occur within the nucleus and that macrophages engulf pyroptotic cellular debris. Furthermore, the expression of the separate ASC domains shows both PYD_A_ and CARD_A_ cluster in filamentous aggregates. PYD_A_ aggregates are sufficient to elicit cell death at a reduced rate, showing CARD_A_ is required both for maximal speck clustering and cell death efficiency. Finally, by generating a Caspase-1 orthologue knockout, we conclude that speck formation unleashes Caspase-dependent pyroptosis in keratinocytes *in vivo*.

## Results

### Tissue specific expression of ASC

ASC has been shown to be expressed in the skin, digestive tract, bone marrow and peripheral blood leukocytes, among other tissues in humans (36), and most myeloid lineage cell lines also express *asc* constitutively (7). However, no encompassing analysis addressing the spatial distribution of its expression sites within an organism has been made. To investigate the role of ASC *in vivo* we first characterized gene and protein expression in zebrafish by Reverse Transcription PCR, *in situ* hybridisation and immunofluorescence with a newly generated antibody against zebrafish ASC. The expression of *asc* is detectable from the morula stage onward, and adult hematopoietic tissues also express *asc* (Fig. S1A). In 3 dpf larvae, *asc* RNA is present throughout the epidermis and in the area around the gills (Fig. 1A and fig. S1B), where it has previously been reported to have a role in pharyngeal arch development (35). Sections showed expression in internal tissues such as the intestinal epithelium and individual *asc*-expressing cells in the brain (Fig. 1B-C and fig. S1C-G”). The lateral line system and some internal tissues, such as the notochord and muscle, lacked ASC. Immunostainings showed ASC’s presence in the epidermis from 1 to at least 5 dpf (Fig. S1I-O). Transgenic tissue-specific markers identified the ASC-expressing cells in the skin as both enveloping layer (EVL) and basal keratinocytes (Fig. 1D and D’ and fig. S1P). In these cells the protein is seen both in the cytoplasm and the nucleus (Fig. 1E and fig. S1P). All macrophages express ASC, as do most neutrophils (Fig. 1F and G), but not all cells labelled by the myeloid lineage reporter *pU1* express ASC (Fig. 1H).

**Fig. 1.**
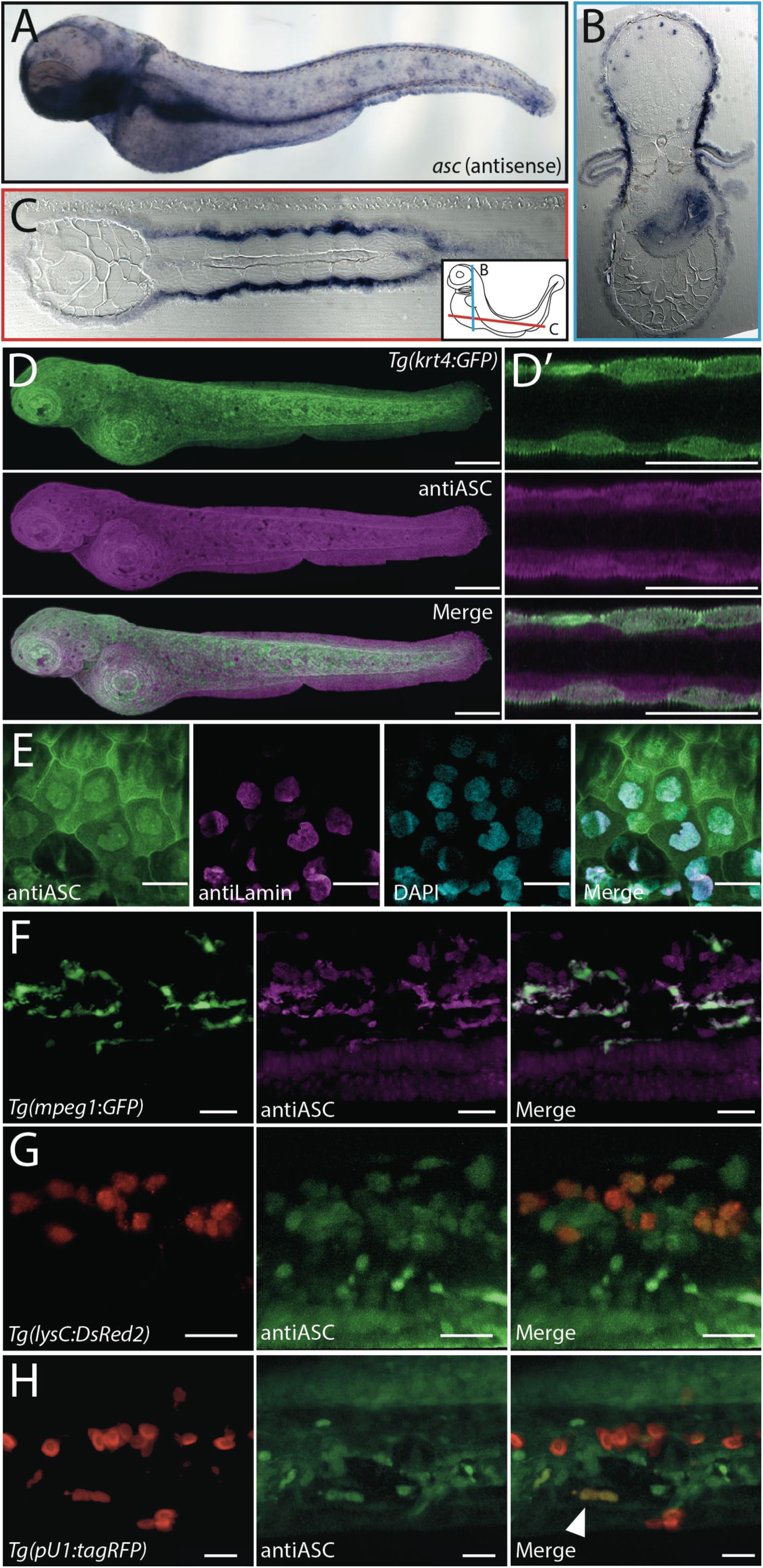
*asc* is expressed during zebrafish early development. *asc* whole-mount *in situ* hybridization (*wish*) of 3 dpf zebrafish larvae [A] with cross [B] and longitudinal [C] sectioning of plastic embedded *wish* sample showing expression in epidermis, intestinal epithelium, and cells located in the brain. Immunostaining of ASC in 3dpf *Tg(krt4:GFP)* larva [D]. Optical cross section of lateral fin showing GFP expression in the enveloping layer (EVL), and ASC expression on both epidermal layers [D’]. Wildtype 3 dpf larva immunostained for ASC, together with nuclear envelope marker Lamin and DAPI shows its nuclear and cytoplasmic localization [E]. Immunostaining of 3 dpf *Tg*(*mpeg:GFP*) [F], *Tg*(*lysC:DsRed2*) [G] and *Tg*(*pU1:tagRFP*) [H] larvae showing expression of ASC in macrophages, neutrophils, and a single myeloid cell in the CHT [H, white arrowhead]. Scale bars, 300 μm for full larvae, otherwise 30 μm.

### Endogenous ASC and specks visualized in vivo in a knockin transgenic line

To be able to study ASC *in vivo* we generated a transgenic CRISPR knockin line through homology-dependent repair in which the endogenous protein is fused with GFP, called *Tg(asc:asc-EGFP)* (Fig. S2A-D). In agreement with the above results, transgenic embryos have ASC-GFP throughout the entire epidermis in nuclear and cytoplasmic compartments as well as in the intestinal epithelium (Fig. 2A-B” and fig. S2E-G). ASC-GFP is also expressed in myeloid cells (Fig. 2C). Microglia, the tissue-resident macrophages of the brain (37) were ASC positive as were cells in the caudal hematopoietic tissue, many (but not all) of which were labelled by the *pU1* reporter transgene. At all stages examined, muscle cells and other internal tissues were devoid of GFP.

**Fig 2.**
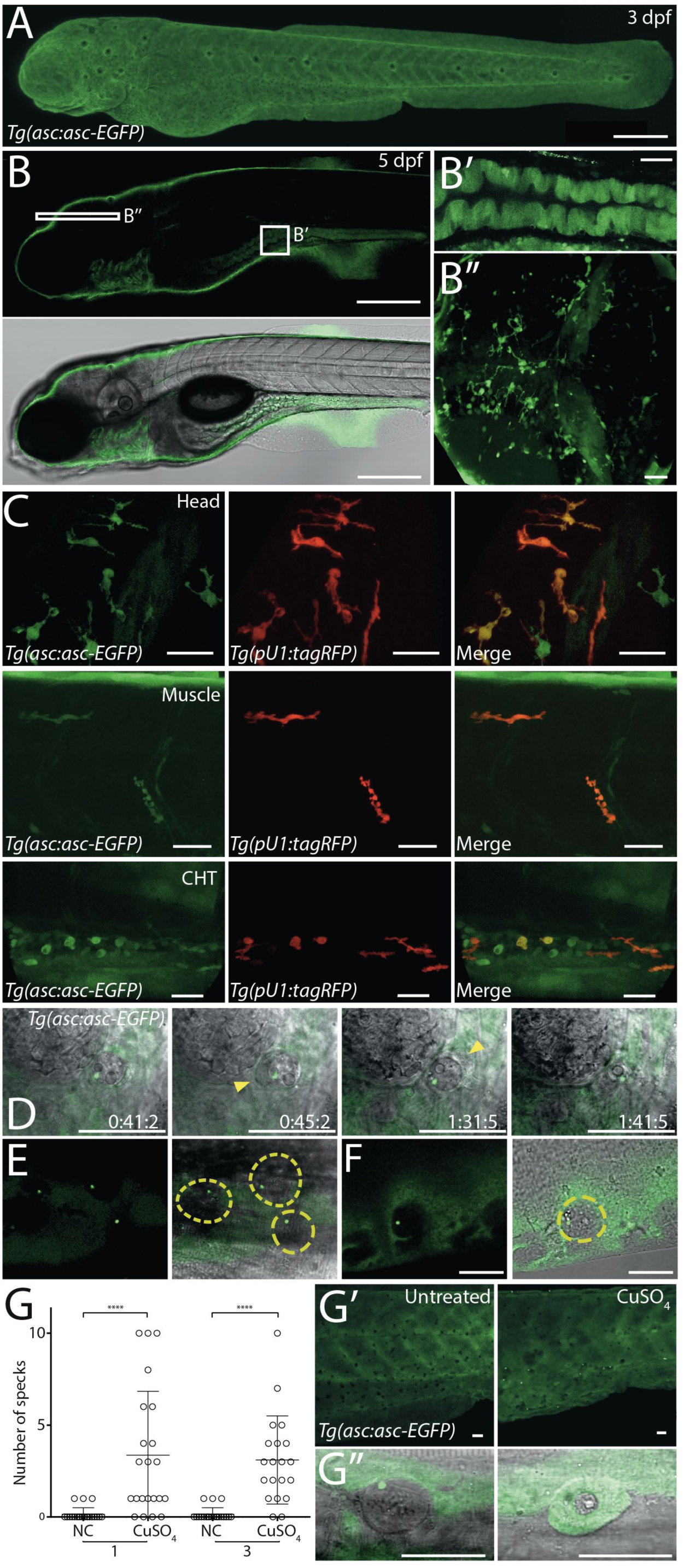
Endogenous ASC forms specks *in vivo* in the *Tg(asc:asc-EGFP)* line. Live imaging of *Tg(asc:asc-EGFP)* 3 dpf [A] and 5 dpf [B] larvae, with intestine [B’] and head [B’’] optical sections. Live imaging of head, muscle and caudal hematopoietic tissue (CHT) of 3 dpf *Tg*(*asc:asc-EGFP, pU1:tagRFP*) larvae [C]. Time lapse imaging of keratinocyte with speck. Single plane merged with the brightfield is shown. Yellow arrowheads highlight a second cell that appears to surround the speck-containing cell [D]. Live imaging of specks in the dorsal epidermis [E] and ventral fin [F] of 3 dpf *Tg(asc:asc-EGFP)* larvae. Merge with brightfield plane shows each speck is within a cell with altered morphology (dashed yellow line). *Tg(asc:asc-EGFP)* 3 dpf larvae were treated with 25 μΜ CuSO_4_ for 1h. At 1 and 3 hours post treatment (hpt) number of specks per larva were quantified (One way ANOVA, \*\*\*\**P*<0.0001) [G]. Live imaging of untreated and treated larvae showing high damage of epidermis and increase in specks [G'], examples in treated embryo of single cells displaying altered morphology with and without speck formation [G’’]. Scale bars, 300 μm for full larvae, otherwise 30 μm.

We observed the sporadic appearance of GFP specks in the epidermis of *Tg(asc:asc-EGFP)* larvae (Movie S1). Without exception, specks were contained in dead or dying cells, as shown in brightfield images where these cells were rounded and dislodged from the rest of the epithelium (Fig. 2D-F). The reason for spontaneous speck formation in these examples is unclear. To determine if inflammatory stimuli could trigger speck formation in epidermal cells, we exposed *Tg(asc:asc-EGFP)* embryos to high concentrations of copper sulphate (CuSO_4_), a compound toxic to zebrafish larvae (38-40). The epidermis of these larvae showed signs of stress, with many deformed cells forming a rugged instead of smooth epithelium, and had significantly increased numbers of specks (Fig. 2G and G’). Cells containing a speck were rounded and dislodged from the rest of the epithelium, which is indicative of cell death. However, not all abnormal epidermal cells had specks (Fig. 2G’’), suggesting CuSO_4_ exposure triggers a range of stress symptoms, and speck formation may occur as an indirect consequence of CuSO_4_-induced toxicity to the skin. Because toxicity-induced speck formation in the skin resulted in undesired side effects, making this an inadequate system in which to address the dynamics and consequences of speck formation *in vivo*, we tested other more direct means of triggering speck formation.

### Speck formation in vivo is induced by NLR or ASC overexpression

When ASC is present at endogenous concentrations, activated members of the NLR protein family, among other receptors, can trigger speck formation. Under overexpression conditions, however, the propensity of ASC to spontaneously aggregate in cultured cells is well documented (11, 18, 19). We therefore tested whether these stimuli resulted in speck formation in live fish. Overexpressing a PYD-containing zebrafish NLR lacking the LRR domain led to ASC-GFP speck formation in epidermal cells of the *Tg(asc:asc-EGFP)* line, showing that the GFP-tagged endogenous ASC responds appropriately to its direct stimulus (Fig. 3A and Movie S2). We also used an overexpression system based on a construct, *HSE:asc-mKate2*, in which mKate2-tagged ASC is expressed under the control of a heat shock promoter (41) that allowed us to induce ASC expression throughout the fish, including cells that do not endogenously express it. Transient expression of ASC-mKate2 from this construct led to the appearance of specks, whereas mKate2 alone had a cytoplasmic distribution (Fig. S3A). Speck formation was not caused by the mKate2 fused to ASC, nor by heat shock-related stress, since overexpressing ASC with other tags and using other expression systems also resulted in speck formation (Fig. S3B-D). To simultaneously and stably induce ASC-mKate2 overexpression in all cells we generated the transgenic line *Tg(HSE:asc-mKate2)* (Fig. 3B). A quantification of speck formation over time in transgenic embryos shows that from 2.5h post heat shock (2.5 hphs) the number of specks increases rapidly and plateaus at around 17 hphs (Fig. 3C and Movie S3). Each cell formed only one speck, concomitant with the depletion of the cytoplasmic pool of ASC-mKate2 (Fig. 3D and Movie S3). Although muscle cells do not express *asc* endogenously, the heat shock-induced ASC-mKate2 also assembled into a single speck in these cells. When we overexpressed ASC-mKate2 in *Tg(asc:asc-EGFP)* embryos, specks that formed in muscle cells were constituted exclusively by ASC-mKate2 (Fig. 3E), whereas in epidermal cells, the endogenous ASC-GFP was recruited to the ASC-mKate2 speck (Fig. 3F and Movie S2). These results suggest that overexpression of ASC or its upstream receptors trigger speck formation and bypass the need for an inflammatory stimulus to activate inflammasome signalling.

**Fig 3.**
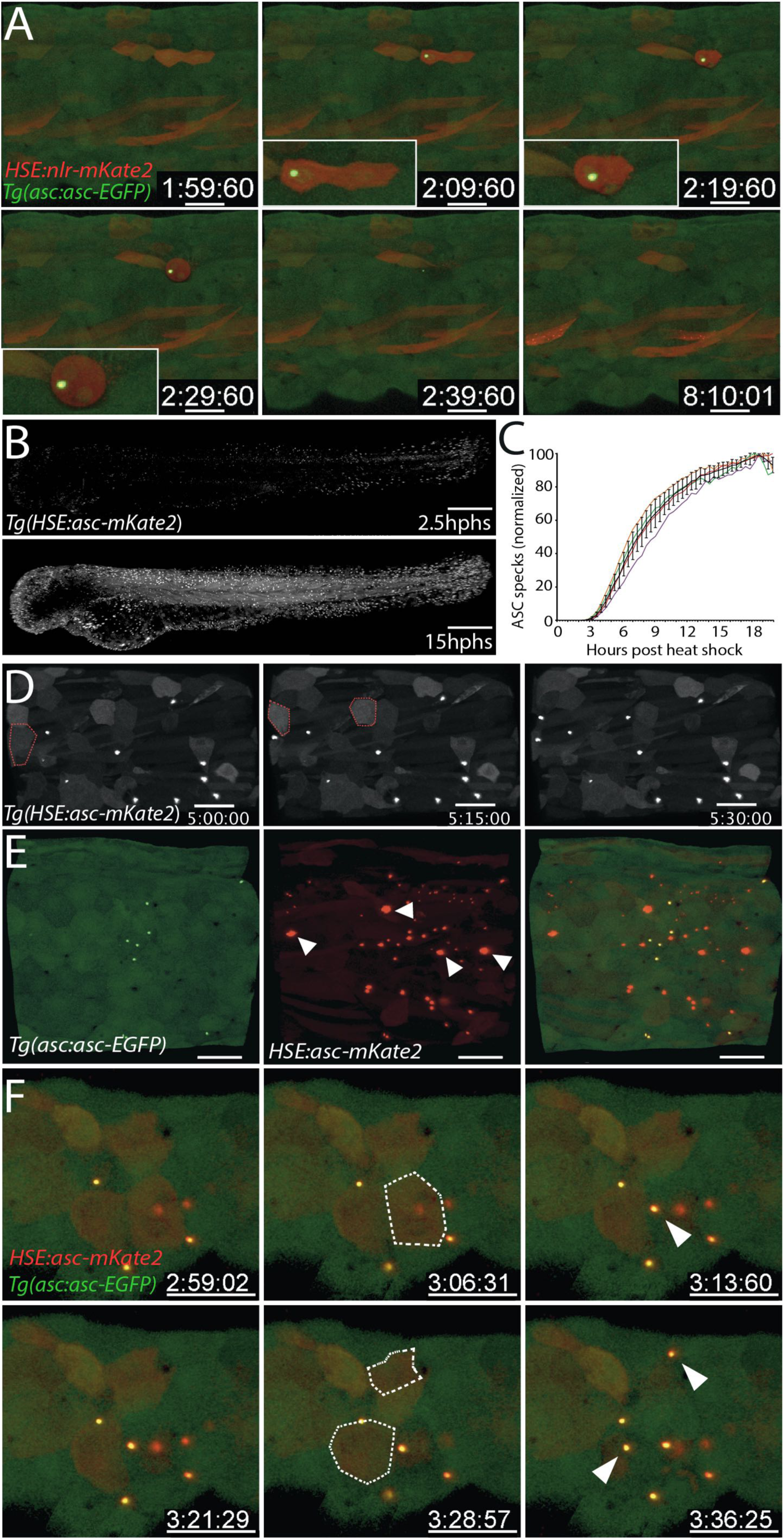
Expression of *asc* or *nlr* induces speck formation. Timelapse imaging of trunk from 3 dpf *Tg*(*asc:asc-gfp*) larva transiently expressing *HSE:nlr-mKate2* at 7 hphs. Inlet shows enlarged view of NLR-mkate2-expressing keratinocytes after speck formation [A]. Time lapse imaging of 3 dpf *Tg(HSE:asc-mKate2)* embryos after heat shock. Shown are timepoints corresponding to 2.5 hphs (upper panel) and 15 hphs (lower panel) [B]. Quantification of speck numbers over entire larvae after heat shock using 3D image analysis software [C]. Time lapse of 3 dpf *Tg(HSE:asc-mKate2)* larvae 3 hphs showing recruitment of ASC-mKate to a single speck per cell (demarcated by dashed red line in timepoint before speck formation). [D]. Live imaging of *Tg*(*asc:asc-gfp*) transiently expressing *HSE:asc-mKate2* 13 hphs [E]. White arrowheads show specks assembled in muscles. Time lapse imaging of *Tg*(*asc:asc-gfp*) larva transiently expressing *HSE:asc-mKate2*. Individual keratinocytes are demarcated with a dashed white line at the time point before the formation of ASC-mKate2 and ASC-GFP double positive specks (white arrowheads). Time lapse was started 3.5 hphs [D]. Single time point of trunk *Tg*(*asc:asc-gfp*) transiently expressing ASC-mKate2. Scale bars, 300 μm for full larvae, otherwise 40 μm.

### Specks are formed by large filamentous assemblies of ASC

Based on cryo-EM structures of *in vitro* assembled PYD_A_ filaments and EM data of ASC specks reconstituted *in vitro* (9), specks are thought to be composed of crosslinked filaments that aggregate into a sphere (42). To characterize the structure of *in vivo*-formed specks, we used correlative light and electron microscopy (CLEM) (Fig. 4A and B, and fig. S4A). We visualized ultrastructural details of specks formed in muscle cells after inducing ASC-mKate2 expression in the *Tg(HSE:asc-mKate2)* line. Specks in muscle cells form a cluster of 700 nm in diameter consisting of highly intercrossed filaments (Fig. 4C, fig. S4B and Movie S4). A three-dimensional model of the filaments reveals that the aggregated ASC filaments form a globular structure (Fig. 4D). This data is a strong indication that the filamentous organization observed from *in vitro* studies is also true of *in vivo* assembled specks.

**Fig 4.**
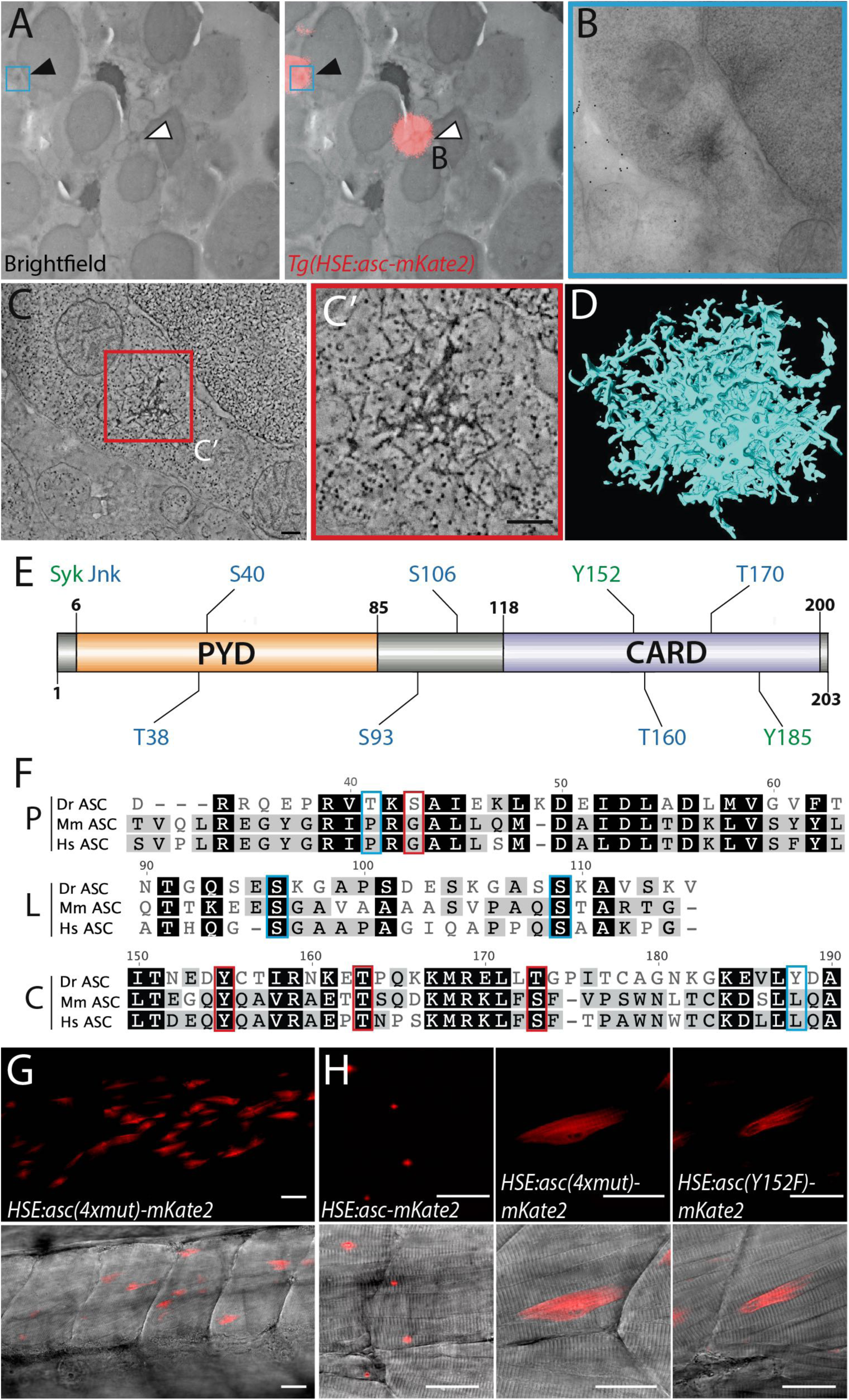
ASC specks are highly intercrossed filamentous structures whose clustering is altered by point mutations. Correlative Light Electron Microscopy (CLEM) of high-pressure frozen 3 dpf *Tg(HSE:asc-mKate2)* larvae 18 hphs [A-D]. Low magnification electron micrograph [A, left panel] and overlay with red channel [A, right panel] imaged with light microscope. White and black arrowheads show location of specks. Area of interest (blue) imaged with electron microscope [B]. TEM tomography slice of speck (black arrowhead) [C] and an enlarged view of intercrossed filaments [C’]. 3D reconstruction of speck after manual tracking of individual filaments [D]. Scale bars, 200 nm. Results from phosphorylation sites analysis using the online tool GPS 2.1.1 depicting Syk and Jnk-specific predicted phosphorylation sites in zebrafish ASC [E]. Portions of zebrafish (Dr), mouse (Mm) and human (Hs) ASC protein alignment separated by domain (P, PYD; L, linker; C, CARD). Aminoacids identified in the analysis are boxed, in red those mutagenized [F]. Live imaging of larvae transiently expressing *HSE:asc(4xmut)-mKate2*, containing 4 missense mutations (T38A, Y152F, T160A and T170A) [G]. Single muscle cell in larvae transiently expressing either *HSE:asc-mKate2* or, *HSE:asc(4xmut)-mKate2* or *HSE:asc(Y152F)-mKate2* [H]. Scale bars, 30 μm.

### Mutating conserved predicted phosphorylation sites abrogates speck formation

Activation of ASC, like other inflammasome components, is subjected to regulation by posttranslational modifications (7). Thus, the speck formation we observe should depend on these as well. We used the overexpression system to test whether ASC in zebrafish was regulated through phosphorylation by the c-Jun N-terminal kinase (Jnk) and spleen tyrosine kinase (Syk) signaling pathways, as reported for mammalian ASC (43, 44). An *in silico* analysis, as used by Hara et al. (2013), predicted a number of potential Jnk and Syk phosphorylation sites in zebrafish ASC (Table S1). Three corresponded to residues within the CARD_A_ that are conserved in mouse and human ASC (Fig. 4E and F). We mutated these three sites (Y152F, T160F and T170A) and one additional site in PYD_A_, which was not conserved (T38A). Since muscle cells do not express *asc* endogenously we were able to use these cells for an *in vivo* analysis of speck formation by mutant proteins while avoiding interference from the wild type ASC. Transiently expressed ASC-mKate2 containing the four mutations formed a striated pattern or large filamentous aggregates in muscle cells, rather than a compact speck (Fig. 4G). By expressing constructs with single mutations, we found that the Y152F mutation was sufficient to disrupt speck formation entirely (Fig. 4H), similar to the corresponding mutations in mouse (Y144A) or human ASC (Y146A) which also caused defective speck formation (43, 44). These results support the notion that speck formation caused by the experimental conditions used here is under the control of conserved ASC post-translational regulatory mechanisms and assembly therefore follows the physiological signalling pathway.

### Speck formation leads to keratinocyte pyroptosis

It is well established that speck formation can cause cell death by pyroptosis in macrophages in culture. However, the first barrier a pathogen must overcome to establish infection are epithelial surfaces that cover the body, which, as we have shown, express high levels of ASC. Yet, very little is known about the function and dynamics of ASC activation and speck formation in this important tissue. Since inducing *asc* expression in the *Tg(HSE:asc-mKate2)* line allows us to study cell-type specific responses to speck formation, we compared responses of keratinocytes, which endogenously express *asc*, to muscle cells, which do not. We observed starkly different responses to speck formation. Keratinocytes round up within minutes after speck formation, whereas muscle cells show no visible change over at least 10 hours, during which the speck continuously increases in size (Fig. 5A and Movie S5). The response in epidermal cells was independent of the method used to overexpress ASC (Fig. S4C). That the appearance of ASC-mKate2 specks is associated with the same morphological changes as those seen after the formation of endogenous ASC-GFP specks suggests that inflammasome signalling is being activated in these cells as a result of overexpression-induced speck formation.

**Fig. 5.**
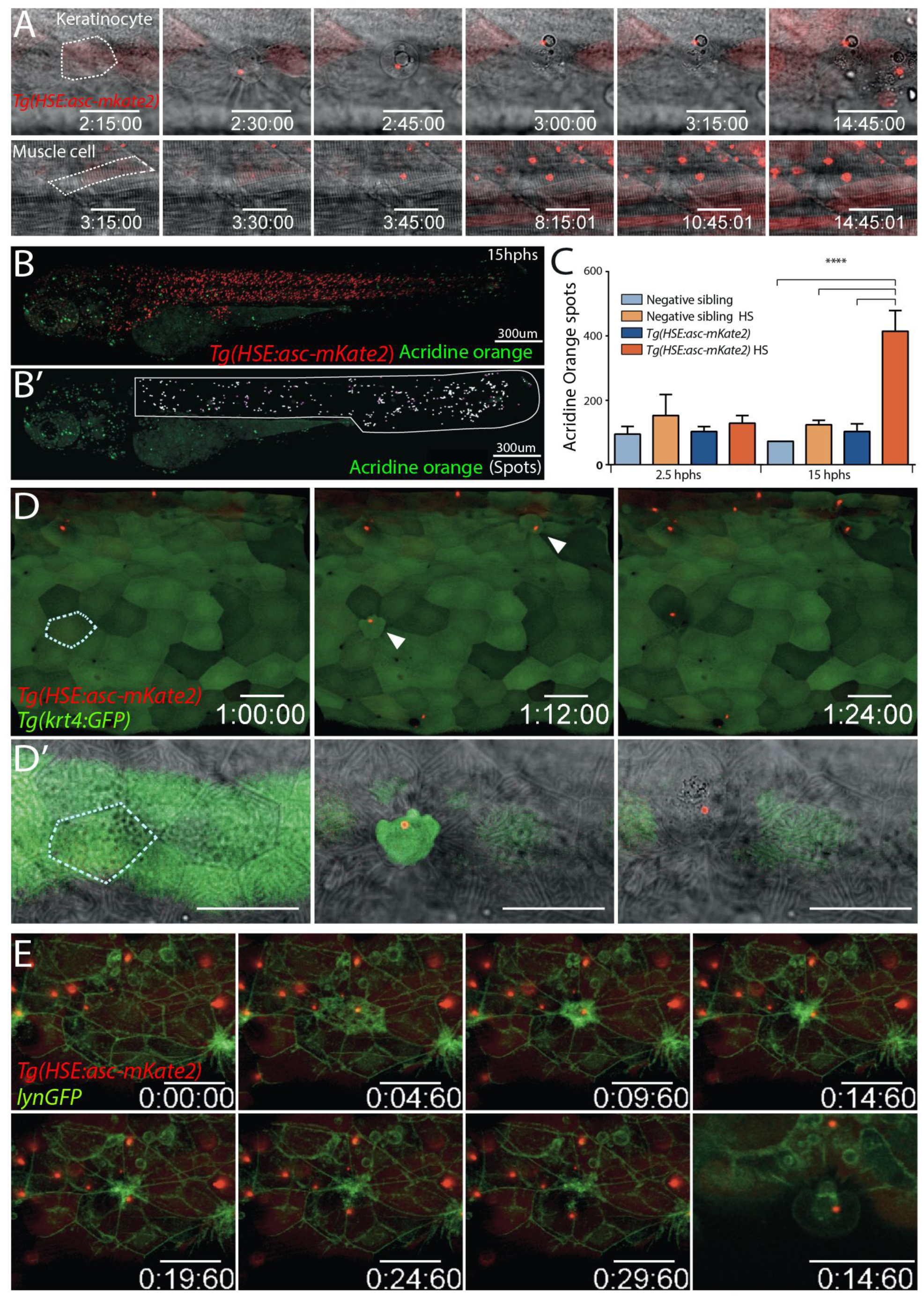
ASC speck formation in keratinocytes leads to cell death. Time lapse imaging of speck formation in keratinocyte (top) and muscle cell (bottom) in 3 dpf *Tg(HSE:asc-mKate2)* larva 3 hphs. Drastic morphological changes occur only in keratinocytes [A]. *Tg(HSE:asc-mKate2)* larvae and negative siblings were stained with acridine orange and imaged 2.5 and 15 hphs [B, upper panel]. 3D rendering of individual larvae manually segmented to exclude the head, heart and yolk regions. Acridine orange spots in segmented region were quantified using 3D image analysis software [white spots, B’]. Spots positive in the red channel were excluded [magenta spots, B’]. Histogram of acridine orange spots in each group shows only transgenic larvae 15 hphs have significantly higher cell death (One way ANOVA, ****P<0.0001) [C]. Time lapse imaging of *Tg(HSE:asc-mKate2, krt4:GFP)* larvae 3 hphs showing morphological changes in EVL keratinocyte upon speck formation [white arrowheads, D, upper row]. Enlarged view of EVL keratinocyte [dashed white outline, D] of single plane with the brightfield [D’]. Time lapse imaging of *Tg(HSE:asc-mKate2)* injected with *lynGFP* mRNA for membrane visualization 8 hphs. Epidermal layer shows extrusion and gap closure after speck formation [E]. Single plane showing extruded keratinocyte [E’]. Scale bars, 30 μm.

We quantified cell death in the *Tg(HSE:asc-mKate2)* line by using acridine orange (Fig. 5B). Before specks assemble, *Tg(HSE:asc-mKate2)* and control larvae show similar levels of staining. However, after speck formation, cell death was significantly higher in heat shocked transgenic larvae (Fig. 5C). Most of the acridine orange staining was located in the skin (Fig. S4D); and keratinocytes, but not muscle cells, accumulated acridine orange in their surroundings after speck formation (Fig. S4E). This, together with the observed changes in morphology, suggested that keratinocytes were undergoing cell death upon speck formation. To test this, we monitored the cellular changes in response to speck formation specifically in EVL keratinocytes using *Tg(krt4:GFP, HSE:asc-mKate2)* larvae (Fig. 5D and Movie S5). All GFP-positive cells that formed a speck showed classic signs of pyroptosis (6) less than 15 minutes after speck formation, including rounding up, detachment from the epithelia and loss of plasma membrane integrity. We analysed the process of cell extrusion by labelling the plasma membrane with a membrane-targeted GFP (lynGFP) and observed that speck formation led to extrusion of the pyroptotic cell from the epithelial sheet, with surrounding cells sealing the gap (Fig. 5E and Movie S5). This was also seen after transient overexpression of ASC-tGFP in a reporter line labelling the membranes of keratinocytes (Fig. S4F and Movie S5). These results show that keratinocytes undergo pyroptosis within 15 min of speck formation.

### Effect of speck formation by nuclear ASC

Both when detected by antibodies and tagged by GFP, endogenous ASC is present in the cytoplasm and the nucleus. Either pool can form specks in HeLa cells (17), although the significance of this, and in particular, whether both nuclear and the cytoplasmic specks can induce cell death *in vivo*, is unclear. To test this, we transiently expressed a nuclear-targeted ASC-mKate2 (NLS-ASC-mKate2) in the *Tg(asc:asc-EGFP)* line, which would allow us to monitor not only the effect of nuclear ASC, but also the endogenous nuclear and cytoplasmic ASC pools. When NLS-ASC-mKate2 formed specks in the nucleus of ASC-GFP expressing keratinocytes, these cells underwent cell death with the same dynamics as described above. Cell death occurred without the recruitment of the cytoplasmic pool of the endogenous ASC-GFP (Fig. 6A and Movie S6). Therefore, the presence of a nuclear speck is sufficient, and neither the depletion of the cytoplasmic pool nor a cytoplasmic speck is required for keratinocyte pyroptosis. However, in cases where the nuclear envelope became permeable to the endogenous ASC-GFP before death occurred, the cytoplasmic pool of ASC-GFP was also recruited to the nuclear speck (Fig. 6B-E).

**Fig 6.**
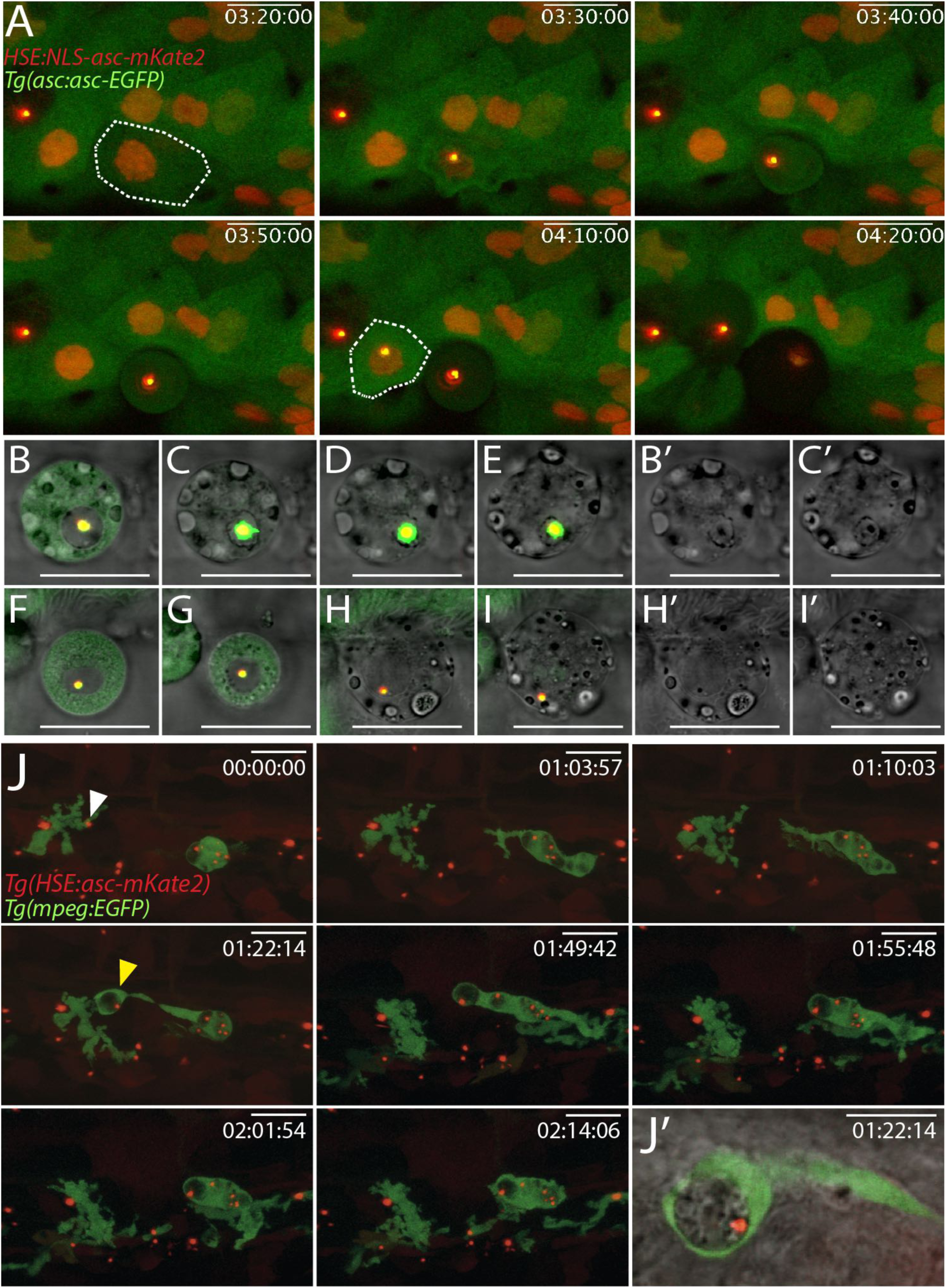
Nuclear specks causing cell death and macrophage engulfment of speck-containing cellular debris. Time lapse of 3 dpf *Tg(asc:asc-EGFP)* larvae transiently expressing *HSE:NLS-asc-mKate2* 6 hphs, showing nuclear speck assembly in keratinocytes (white dashed line) leads to cell death [A]. Cell undergoing cell death with nuclear speck and without depletion of ASC-GFP in the cytoplasm [B-E]. Brightfield of respective timepoints show breakdown of nuclear envelope allows recruitment of cytoplasmic ASC-GFP [B’ and C’]. Loss of plasma membrane integrity [F-I] prior to nuclear envelope breakdown, results in leakage of cytoplasmic ASC-GFP as shown in brightfield [H’ and I’]. Time lapse imaging of *Tg(HSE:asc-mKate2, mpeg:EGFP)* larva 17 hphs shows macrophage engulfing a speck (white arrowhead) [J]. Brightfield merge of single plane showing phagocytic cup (yellow arrowhead) [J’]. Scale bars, 20 μm.

In cases where the plasma membrane collapsed prior to nuclear envelope breakdown, cytoplasmic ASC-GFP leaked to the extracellular environment before it was recruited to the nuclear speck (Fig. 6F-I). Similar results were obtained by transiently coexpressing ASC-mKate2 with GFP in a transgenic line carrying the *βactinNLS-tagBFP* transgene to label all nuclei (Fig. S5 and Movie S6). Namely, specks assembled either from the cytoplasmic or the nuclear pool of ASC, but regardless of the compartment in which the speck formed its assembly led to cell death. This confirms that speck formation in the nucleus is sufficient to trigger pyroptosis in keratinocytes.

### Clearance of pyroptotic debris containing ASC specks by macrophages

After macrophages undergo pyroptosis, they leave behind a structure composed of ruptured plasma membrane containing insoluble contents called “pore-induced intracellular traps (PITs)”. In culture, neighbouring phagocytes clear up PITs through efferocytosis (45). There is also evidence that ASC specks are released to the extracellular space and can spread inflammation by recruiting the soluble ASC in the cytoplasm of phagocytes that engulf them (46, 47). However, whether ASC specks remain trapped in PITs, and the rules that determine when engulfed specks induce speck formation and pyroptosis in the phagocyte have yet to be defined. We observed that, after keratinocyte cell death, specks remained enclosed within the cellular debris (Fig. 2E and F and fig. 5A). To test whether phagocytes could engulf speck-containing cellular debris, we induced ASC-mKate2 expression in the *Tg(HSE:asc-mKate2)* line crossed with the macrophage reporter line. Macrophages were indeed capable of engulfing pyroptotic debris with specks (Fig. 6J and Movie S7). Instances of macrophages containing multiple phagosomes with specks suggest there is continuous uptake of speck-containing debris by phagocytes, and that engulfed specks do not elicit a pyroptotic response in the macrophages within 2-3 hours after engulfment. Instead, the gradual loss of fluorescence from phagocytized ASC-mKate2 specks suggests that macrophages are capable of digesting specks after engulfment (Fig. S6 and Movie S7). Thus, the main function of phagocytes that we observe *in vivo* is to clear speck-containing pyroptotic cellular debris, and we have seen no incidences of specks triggering further death after engulfment.

### Domain requirements for compact speck clustering and efficient cell death

Based on *in vitro* and cell culture experiments, the PYD and CARD domains of ASC are thought to have distinct roles during speck formation, with PYD_A_ assembling into filaments that are crosslinked by inter-filament CARD interactions (15). To determine each domain’s role in speck assembly and pyroptosis *in vivo* we overexpressed the single PYD_A_ and CARD_A_ fused to mKate2 (PYD_A_-mKate2 and CARD_A_-mKate2, respectively). In muscle cells, PYD_A_ most frequently assembled into long filamentous structures, whereas CARD_A_ aggregated into smaller punctate aggregates throughout the cell (Fig. 7A). In contrast, expression of either domain keratinocytes resulted in the formation of a normal-looking compact speck that led to pyroptosis (Fig. S7A and B, and Movie S8). The most likely reason for this difference is the presence of endogenous ASC in keratinocytes. To test this, we repeated these experiments under conditions of *asc* morpholino knockdown (Fig. S7C). While overexpressed ASC*-mKate2 under *asc* knockdown conditions formed compact specks in keratinocytes and caused cell death (Fig. S7D and Movie S8), as observed in control larvae, overexpressed PYD*_A_ or CARD_A_ failed to do so. Instead, following a slower depletion of the cytoplasmic pool of the protein than that of full-length ASC, the single domains formed aggregates similar to those assembled in muscle cells (Fig. 7B and C and Movie S8). The formation of these aggregates was not associated with immediate cell death: PYD_A_- expressing epidermal cells died over 2 hours after PYDA-aggregates are first seen whereas cells with CARD_A_ aggregates survived for more than 10 hours after aggregate formation. This differs from the fast response observed within ~10 min of ASC-mKate2 speck formation in *asc* knockdown larvae. PYD_A_ is therefore both necessary and sufficient for cell death, which suggests that this domain mediates the interaction with downstream elements that trigger pyroptosis.

**Fig 7.**
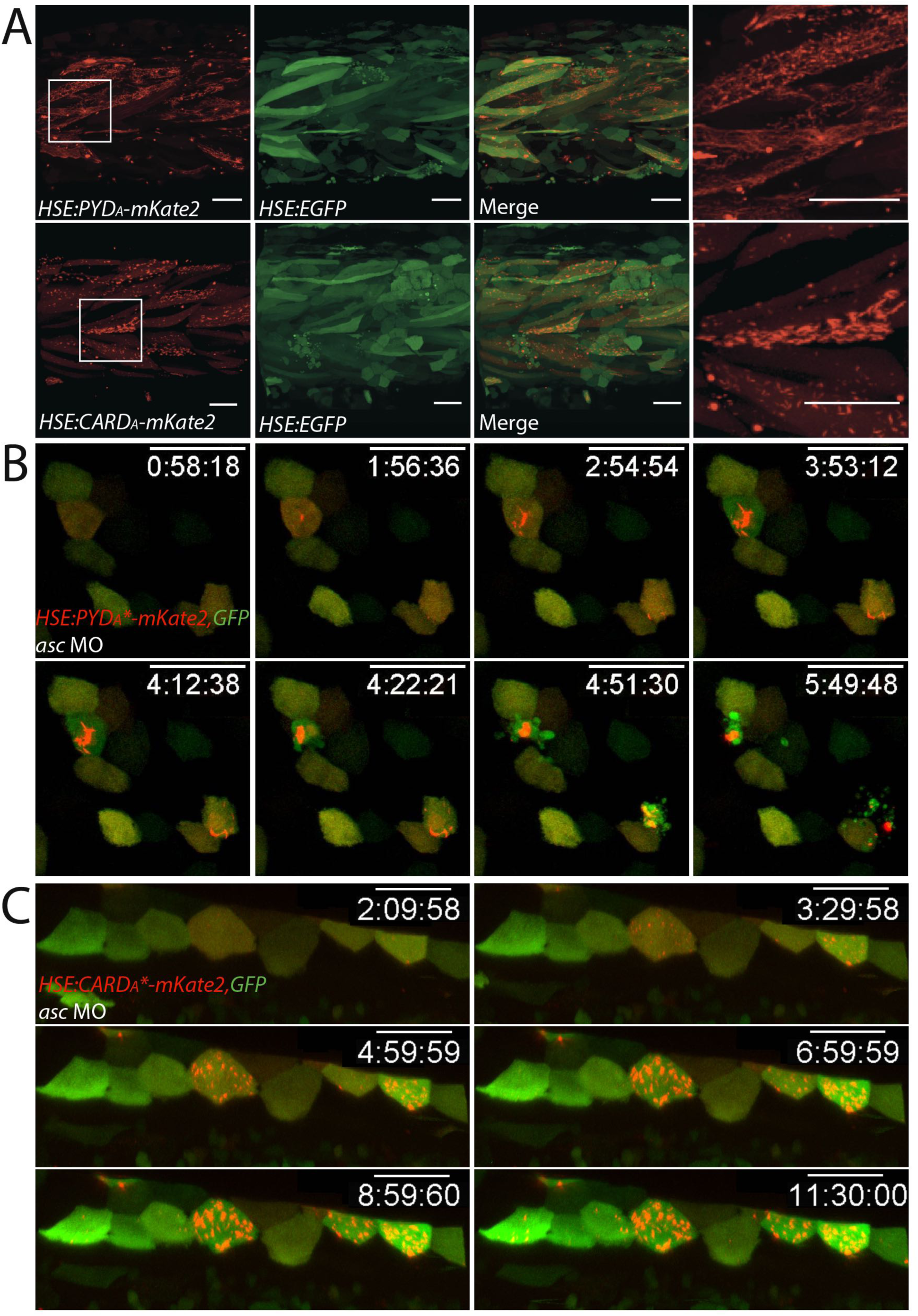
PYD aggregates lead to delayed pyroptosis. Live imaging of 3 dpf larvae transiently expressed *HSE:PYD_A_-mKate2* or *HSE:CARD_A_-mKate2* with GFP 17 hphs [A]. Expression of either domain leads to the formation of filamentous aggregates of varying lengths in muscle cells. Time lapse imaging of *asc* morpholino-injected *Tg(asc:asc-EGFP)* larvae transiently expressing the *asc* morpholino resistant *HSE:PYD*_A_-mKate2* [B] or *HSE:CARD_A_-mKate2* [C] with GFP. If endogenous ASC is absent, PYD_A_ aggregates cause cell death 2 hours after the aggregates first form, whereas CARD_A_ aggregates do not, even 10 hours after their assembly. Scale bars, 40 μm.

### PYD-dependent recruitment of Caspa to the ASC speck

In mammals, the effector domain of ASC for triggering pyroptosis is the CARD, which interacts with the CARD of Caspase1. For this reason, it is surprising that in zebrafish PYD appears to be the effector domain. We therefore tested whether caspases were involved in the response to speck formation, and if so, how they interacted with ASC. Treatment of *Tg(HSE:asc-mKate2)* larvae with the pan-caspase inhibitor (Q-VD-OPh hydrate) resulted in a significant reduction in cell death without affecting speck formation (Fig. 8A and B), showing that caspase activity is required for ASC-dependent pyroptosis. Since caspases are recruited to the speck for auto-activation (4), we tested which caspases could interact with the ASC speck. There are two homologues of mammalian *caspase-1* in zebrafish, *caspa* and *caspb*, both with N-terminal PYD domains. We generated GFP fusions for both caspases, as well as for *casp3a*, the zebrafish orthologue of mammalian Caspase-3, and transiently coexpressed them with ASC-mKate2. Only Caspa was recruited to ASC specks assembled in muscle cells (Fig. 8C). By expressing the PYD and p20-p10 domains of Caspa (PYD_C_ and p20-p10) separately with either the PYD_A_ or CARD_A_, we confirmed that the interaction occurs via the PYD domains of both proteins (Fig. 8D and fig. S8A-C).

**Fig. 8.**
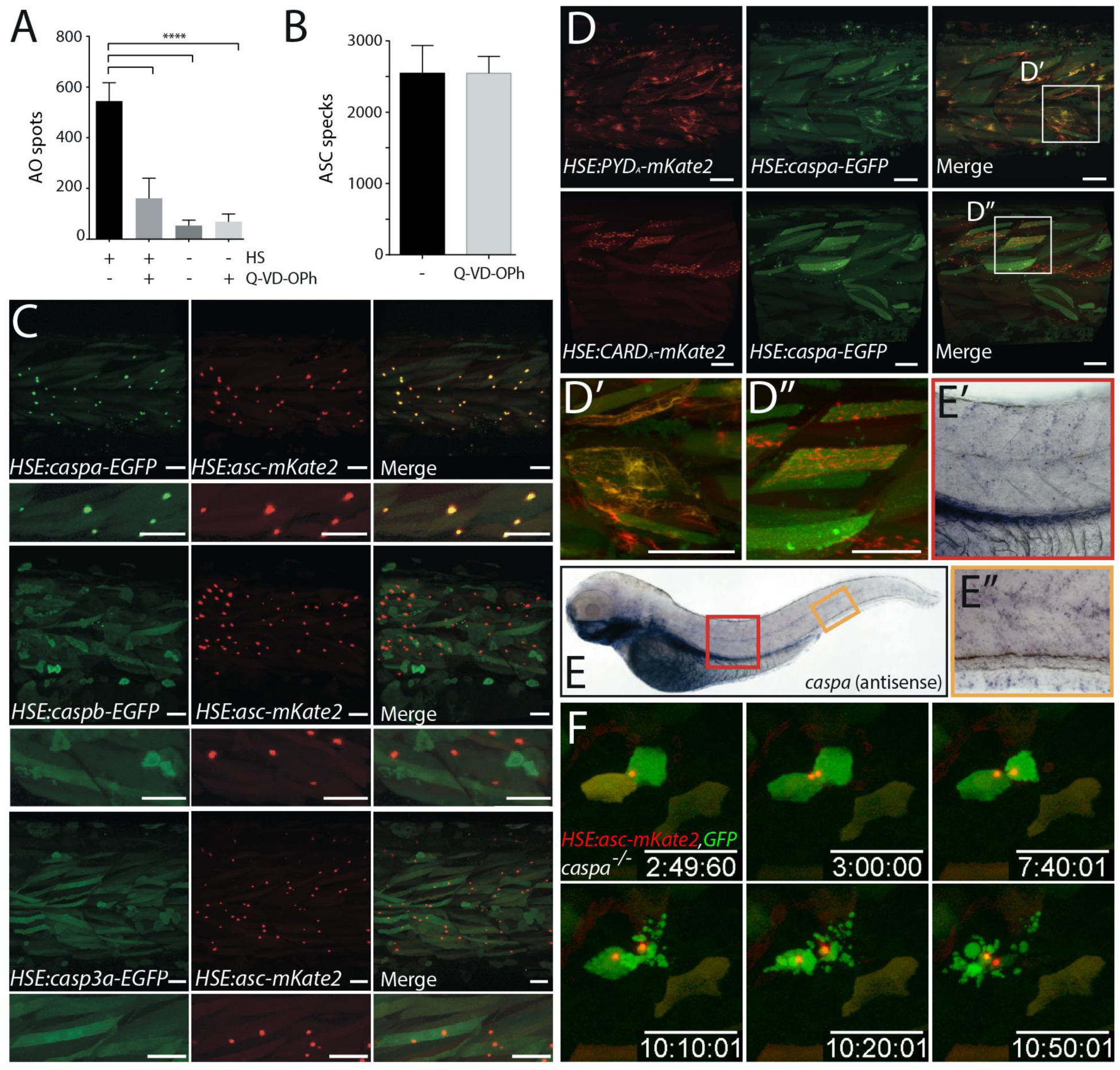
ASC speck formation leads to pyroptosis via activation of Caspa through PYD-PYD domain interaction. 3 dpf *Tg(HSE:asc-mKate2)* larvae treated with the pan-caspase inhibitor Q-VD-Oph (100 μΜ) after or without heat shock were stained with acridine orange at 17 hphs. Acridine orange spots [A] and specks [B] were quantified. Treatment with Q-VD-Oph significantly diminished cell death caused by speck formation compared to non-treated controls (One way ANOVA, \*\*\*\**P*<0.0001). Live imaging of transient expression of *HSE:caspa-EGFP*, *HSE:caspb-EGFP* or *HSE:casp3a-EGFP* with *HSE:asc-mKate2* [C]. Recruitment to the ASC-mKate2 specks only occurs in the case of Caspa-GFP coexpression. Live imaging of heat-shock induced transient expression of *HSE:PYD_A_-mKate2* or *HSE:CARD_A_-mKate2* with *HSE:caspa-EGFP* in 3 dpf larvae 19 hphs [D] with enlarged view of single cells [D' and D'']. PYD_A_, but not CARD_A_, aggregates recruit Caspa-GFP. *caspa* antisense *wish* in 3 dpf larvae [E]. Enlarged view shows expression in skin and ventral fin [E' and E'']. Time lapse imaging of *caspa* mutants transiently expressing *HSE:asc-mKate2* with GFP 3 hphs [F]. Cell death response is severely affected in *caspa^-/-^* keratinocytes, with cells dying an apoptotic-like death more than 7 hours after speck formation. Scale bars, 40 μm.

Transient overexpression of Caspa, unlike that of Caspb or Casp3a, was extremely toxic to epidermal cells (Fig. S8D). Caspa-GFP-overexpressing embryos lacked normal-looking keratinocytes with homogeneous GFP expression, and instead had copious green-labelled cellular debris. Even muscle cells, which were not affected by ASC speck formation, displayed signs of damage after Caspa expression (Fig. S8E). Considering that endogenous *caspa* is expressed in the skin (Fig. 8E and fig. S8F) these data strongly suggest that Caspa is the effector caspase activating pyroptosis in keratinocytes after speck formation and that muscle cells are protected from speck-induced pyroptosis because they do not express it.

To test this hypothesis, we generated a *caspa* mutant by use of CRISPR/Cas9 and identified two mutations (*caspa*^K^** and *caspa*^Δ800^) that resulted in transcripts with a nonsense codon within the first exon (Fig. S8G-J). We transiently expressed ASC-mKate2 and GFP in *caspa* knockout larvae. Speck formation in keratinocytes proceeded normally in these larvae, but did not result in pyroptosis with cells instead surviving for hours after speck formation (Fig. 8F and Movie S9). Eventually, keratinocytes with specks displayed cellular blebbing, nuclear condensation and slowly disintegrated into vesicles strongly reminiscent of apoptotic bodies, suggesting that if Caspa is absent, speck formation results in activation of apoptosis instead of pyroptosis. These results establish Caspa as the direct and only downstream effector of ASC speck formation driving immediate pyroptosis *in vivo*.

## Discussion

ASC speck formation is a hallmark of inflammasome activation. The use of cell lines has significantly contributed to dissect the molecular interactions involved in this signalling cascade but we lack deeper understanding of how inflammasome activation occurs in cells within their native environment. This knowledge gap can be bridged by using models that enable visualization of immune processes in the context of the whole organism (27, 30). Previous studies had suggested that some elements of the inflammasome signalling cascade are involved in the defence against pathogens using zebrafish infection models (*48-50*) and it was recently shown that zebrafish lacking ASC are more susceptible to *Salmonella* Typhimurium infection (51). In our case, a live imaging approach allowed us to characterize inflammasome signalling in the skin *in vivo*. In both fish and mammals, the skin functions as an immune organ that provides a crucial protective barrier (52). Keratinocytes both relay environmental signals to immune cells and execute a response themselves, with their death acting as a potent trigger of skin inflammation (23, 53). Inflammasome activation in keratinocytes has been implicated in response to a number of stimuli (25, 54-58) and the strong expression of ASC we observe in the skin, as well as other epithelia like gills and intestine suggested that the activation of inflammasome is of particular importance in these tissues. Our finding that keratinocytes respond to inflammatory conditions by forming ASC specks and triggering pyroptosis, underscores the relevance of inflammasome signalling in epithelia *in vivo*.

Our work shows that the specific structural mechanisms that lead to ASC’s assembly into specks are conserved between zebrafish and mammals. First, several different ways of overexpressing ASC *in vivo* confirm its high tendency for aggregation, consistent with previous examples showing zebrafish ASC specks in mammalian cells (35) and in uninfected control zebrafish larvae injected with *asc-GFP* mRNA (49). Second, the abrogation of speck formation when predicted conserved phosphorylation sites of zebrafish ASC are mutated suggests conservation of Jnk and Syk-dependent posttranslational regulatory mechanisms of ASC (43, 44). Lastly, our CLEM analysis, which constitutes the first structural analysis of *in vivo* specks, shows its clustered filamentous nature and confirms the model based on *in vitro* inflammasome reconstitutions depicting a speck as a three-dimensional globular ultrastructure composed of multiple highly intercrossed filaments (9).

An important difference between mammalian and zebrafish ASC is the domain that interacts with the effector caspase. In contrast to the mammalian inflammasome, in which Caspase-1 and ASC interact via their CARD domains, zebrafish Caspa, which has an N-terminal PYD instead of a CARD, is recruited to the ASC speck via its PYD domain, in agreement with previous mammalian cell culture experiments (35). CARD_A_ in mammals is located on the surface of ASC filaments, enabling the recruitment of Caspase-1. Since CARD domains can themselves assemble into filaments, as in the case of MAVS in RIG-I antiviral signalling (59) the ASC filament domain structure may be inverted in zebrafish, allowing the PYD to interact with Caspa.

Our results on the effects of expressing the individual domains of ASC reveal a correlation between the compaction of the ASC speck and the efficiency with which it leads to cell death. Both PYD_A_ and CARD_A_ alone have the capacity to aggregate when overexpressed, but neither cluster in a single compact speck. CARD_A_ aggregates have no detrimental effect on cells, but overexpression of only PYD_A_ whose aggregates are able recruit Caspa, results in cell death. Therefore, in this setup, neither the association of CARD and PYD, nor the formation of a compact speck, nor the bridging of PYD to other molecules via CARD, are essential for cell death as such. Instead, it appears that the PYD-mediated recruitment of Caspa is sufficient. However, the finding that the rate of aggregation and cell death are significantly reduced indicates that CARD_A_ is needed for the highly efficient and rapid triggering of pyroptosis. This could be achieved by maximizing speck compaction though filament crosslinking, as shown in cell culture (15,16), which might cause more rapid and efficient nucleation and clustering of Caspa than PYD aggregates, by recruiting additional accessory molecules to the speck that accelerate Caspa activation, or through a combination of both mechanisms.

Specks had been shown to remain as stable aggregates in the extracellular space after ASC overexpression in COS-7 cells and in the supernatant of macrophage cell cultures upon exposure to inflammasome-activating stimuli (46, 47, 60). In the *Tg(HSE:asc-mKate2)* line, ASC specks persist after the death of the cells and appear to remain associated with the pyroptotic cellular debris, which can be readily engulfed by macrophages, as is the case in culture for *in vitro* assembled specks (47) and PITs (45). Macrophages *in vivo* continuously cleared up speck-containing cellular debris, and a single macrophage could contain multiple phagosomes with specks. Furthermore, engulfment led to the degradation of the specks within phagosomes.

Franklin et al. (2014) reported that macrophages that engulfed *in vitro* assembled specks could undergo pyroptosis after the speck was released into the cytosol and nucleated clustering of the phagocytes’ soluble ASC (47), an observation which is supported by recent *in vivo* data (61). However, we did not find that a macrophage’s ability to clear up debris *in vivo* diminished or that the macrophage was affected by the engulfment of a speck short term. It is possible that specks enclosed within ruptured membranes are less efficient triggers of the phagolysosomal damage that releases them into the cytosol; or that, *in vivo*, additional conditions are required to activate this mechanism of signalling spreading, such as extraordinarily high or sustained organismal inflammation levels. This would explain why extracellular specks are detected in the case of chronic, but not acute inflammation (47).

We noticed that not all specks that form in the epidermis are removed. Keratinocytes belonging to the outer epidermal layer (EVL), marked by the *krt4* transgene, are extruded from the epithelium towards the outside of the body. Since they are sloughed off and become separate from the living tissue, macrophages are likely unable to reach and remove their cellular debris.

Recently, speck formation within a tissue was visualized by intravital imaging of macrophages derived from retrovirally transduced ASC-GFP hematopoietic stem cells in bone marrow chimeric mice (61). A second study generated a transgenic mouse carrying ASC-citrine that can be expressed in a lineage-specific manner (21). Although both studies analyse inflammasome activation within a living tissue, they rely on the insertion of an additional copy of ASC-FP expressed under viral promoters for protein visualization; thus, expression levels from the transgene are artificial, and cells that endogenously express *asc* will therefore have an increased concentration of the protein. These disadvantages are circumvented by endogenous tagging of *asc*, as in the *Tg(asc:asc-EGFP)* line, in which ASC-GFP is only present in cells where it is endogenously expressed and at physiological levels, thus avoiding activation artefacts. We cannot entirely exclude that the GFP itself influences the behaviour of the protein, but this would be a caveat affecting all studies using fluorescent proteins to visualize ASC live. However, since endogenous inflammasome activation in the context of organismal infection has not been studied live, we believe that the *Tg(asc:asc-EGFP)* line will prove a valuable tool to address this question *in vivo*.

## Experimental Procedures

### Imaging

For confocal microscopy, larvae were anesthetized with MESAB (ethyl-m-aminobenzoate methanesulfonate) by adding the compound to the media at a concentration of 40 μg/ml and mounted in 1.3% low-melting point agarose (Peqlab). Imaging of immunostainings was carried out in a Leica SP8 TCS confocal microscope using dry 20x/0.8 or water 40x/1.1 objectives. Live imaging was performed using Zeiss LSM 780 confocal microscope at room temperature. For time-lapse imaging of epidermal and muscle cells, a 40x water objective was used (LD C-Apochromat 40x/1.1 W Corr M27 or C-Apochromat 40x/1.2 W Corr M27, Zeiss). Whole larvae were imaged using a 5x (Plan-Apochromat 5x/0.16 M27, Zeiss) or 10x (Plan-Apochromat 10x/0.45 M27, Zeiss) as tiles and later stitched.

### asc knockdown

Design and synthesis of *asc* ATG morpholino (5’-GCTGCTCCTTGAAAGATTCCGCCAT-3’) was carried out by Gene Tools, LLC. Stock morpholino was and diluted in nuclease-free H_2_O to a concentration of 3 mM and stored at room temperature. For knockdown experiments, morpholino was injected at a concentration of 0.6 mM. Morpholino was validated by immunostaining and, for *in vivo* experiments, by loss of fluorescence after injection in homozygous *Tg(asc:asc-EGFP)* embryos.

### Generation of asc:asc-EGFP line

*sgRNA design:* Guide RNAs that targeted the last exon of *asc* (ENSDARG00000040076) were designed using the CRISPR/Cas9 target online predictor CCTop (http://crispr.cos.uni-heidelberg.de) (62). Two suitable hits, Guide 1 (ATTCCTGATGGATGACCTTG) and Guide 2 (ATCTTCACTCAGCATCCTCA) were synthetized using the oligo annealing method into vector DR274. DR274 was a gift from Keith Joung (Addgene plasmid #42250) (63). *sgRNA in vivo validation:* to test whether sgRNAs Guide 1 and 2 targeted the region of interest *in vivo*, they were individually injected in varying concentrations (15-150 ng/μl) together with 1 μl of Cas9 protein (4 mg/ml) complemented with ca. 150 mM KCl into fertilized eggs at the one-cell stage of the zebrafish TLF strain. Successful knockdown was verified by sequencing of a 1.3 kb PCR product from the targeted region of *asc* (Fwd: CCTGTCTGACCATGTGAACATCTA, Rev: TTAGCATTTGTCCTTATCGCAAAC). *Donor vector construction:* Donor vectors were constructed via Golden GATEway cloning (64). In short, 50 ng of entry vector (EV) plasmids numbered 1 to 6 and a vector backbone, were digested with 0.5 μl of *Bsal* (Fast Digest, Thermo Fisher Scientific) and ligated with 0.5 μl of T4 DNA Ligase (30 U/μl, Thermo Fisher Scientific) in several rounds in one continuous reaction of 10 cycles consisting of 30 min at 37°C and 20 min at 16°C, followed by 5 min of 50°C and 5 min of 80°C to inactivate both enzymes. EV1 included a donor plasmid specific target site for *in vivo* plasmid linearization (GGCGAGGGCGATGCCACCTACGG) (62), EV3 contained an *EGFP* CDS with a flexilinker for tagging of *asc*, EV4 was empty and EV6 contained a STOP codon. Homology 5’ and 3’ flanks of different lengths (1 kb for 5’ and 1 kb or 2 kb for 3’) were amplified from zebrafish gDNA and cloned into empty EV2 and EV5. Flanks were amplified and designed according to the specific Cas9 cleavage sites for Guide 1 and Guide 2 as previously reported (65) to increase chances of precise integration. All vectors whose cloning is not mentioned were kindly provided by the Wittbrodt lab. *Injection:* For homologous recombination, the *asc* sgRNA Guide 1 or 2 (120 ng/μl) and a corresponding donor vector (20-50 ng/μl) were injected with a donor specific sgRNA for donor *in vivo* plasmid linearization (150 ng/μl) and 1 μl of Cas9 protein (4 mg/ml) in a solution complemented with ca. 150 mM KCl. *Screening:* Larvae were screened at 2 dpf for GFP expression. We observed higher successful recombination rates when using *asc* Guide 2 and a donor vector with 5’ and 3’ homology flanks of 1 and 2 kb, respectively. However, the number of positive embryos was low and highly variable, ranging from 1 in 40 to 1 in 200 injected embryos. In total, 18 positive F0 larvae were raised into adulthood and screened for positive integration in the germline by outcrossing with wild type fish. One founder whose F1 progeny carried an allele with a correct insertion of *linker-EGFP* cassette at a rate of 30% was found. Successful integration was confirmed by amplification of the targeted region in the *asc* locus by PCR and sequencing (SF4). Heterozygous *asc-EGFP/+* embryos were raised and incrossed to obtain homozygous *asc-EGFP* embryos.

### Chemical and inflammatory treatments

*Caspase inhibition:* The pan-caspase inhibitor Q-VD-OPh hydrate (Sigma-Aldrich) was resuspended in DMSO at a stock concentration of 10 mM. For caspase inhibition the compound was added directly to the medium at a concentration of 100μΜ. *CuSO_4_ treatment:* 3 dpf larvae were treated with for 1h with Copper (II) sulphate (Sigma-Aldrich) at 25 μΜ. The compound was washed off and specks were quantified 1 or 3 hours post-treatment.

## Acknowledgements

We thank B. Bajoghli for helpful discussion and S. Kraus for technical assistance. We are grateful to F. Peri for zebrafish hosting and J. N. Buffoni and C. Henkel for caretaking, to M. Hammerschmidt for the sharing of the *Tg(krt19:dTomato-CAAX)* and *Tg(krt4:GFP)* zebrafish lines and to D. Gilmour for the *Tg(βactin:NLS-tagBFP)* line generated by L. Newton. We thank the EMBL Protein Expression and Purification Facility as well as the EMBL Animal House for their role in generating the zebrafish ASC antibody, the EMBL Advanced Light Microscopy Facility (ALMF) for continuous support and Zeiss for support of the AMLF. We are grateful to D. Gilmour, A. Meijer and F. Peri for comments on this manuscript.

## Author contributions

P.K. and M.L. designed the study.

P.K. generated the *Tg(HSE:asc-mKate2)*, *Tg(asc:asc-EGFP)* and *caspa^-/-^* lines and performed all experiments except CLEM, which were performed with N.S. and Y.S. T.T. and J.W. contributed with initial *Tg(asc:asc-EGFP)* design. P.K. and M.L. interpreted the data and wrote the paper. All authors read and edited the manuscript.

## Supplemental Information

### Supplemental Experimental Procedures

#### Zebrafish care, transgenic lines and genotyping

Zebrafish (*Danio rerio)* were cared for as described previously (1). The chemical 1-phenyl-2- thiourea (PTU, Sigma-Aldrich) was added to E3 medium at a concentration of 0.2 mM to inhibit pigmentation. The Tupfel Long Fin (TLF) strain was used as wild type. The following transgenic lines were used: *mpeg1:EGFP* (2), *pu1:Gal4-UAS-TagRFP* (3), *lysC:DsRed2* (4), *βactin:ΝLS-tagBFP* (Lionel Newton, unpublished), *krt4:GFP* and *krt19:Tomato-CAAX*(5). Lines generated in this study are described below. gDNA was extracted from full larvae or adult fin clips using the QuickExtract DNA Extraction Solution (Epicentre), genotyping was carried out with Phusion High-Fidelity DNA Polymerase (Thermo Fisher Scientific). All animal experiments described in the present study were conducted under the rules of the European Molecular Biology Laboratory and the guidelines of the European Commission, Directive 2010/63/EU.

#### Acridine orange staining

Acridine orange is a live dye that has previously been used to label dying cells in live zebrafish embryos (6). Larvae were stained by immersion for 45 min in a 1:1500 dilution of a 10mg/ml stock (Sigma-Aldrich) prepared in E3, rinsed to remove excess dye, anesthetized, mounted and imaged directly afterwards. Because the dye is light-sensitive, larvae were kept in the dark during staining.

#### Cloning of expression vectors and expression induction

All expression vectors were coinjected with transposase mRNA (100 ng/μl) in embryos at onecell stage. For all heat shock-driven expression, the fusion protein of interest was cloned into a vector backbone containing a bidirectional heat shock element (HSE) as promoter (7), Tol2 sites for transgenesis and carrying the *cmlc2:tagRFP* as a transgenic marker (8). To induce expression, injected embryos with red “bleeding heart” expression were heat-shocked at 39 °C in a heating block at any stage between 2.5 dpf and 3.5 dpf. The transgenic *HSE:asc-mKate2* line was generated by raising embryos (F0) carrying the heart marker without exposing them to heat-shock. The *ubi:LexPR,LexOP:asc-mKate2* vector containing the LexPR/LexOP transactivation system (9) was generated via Gateway recombination cloning (Thermo Fisher Scientific) of *ubi(p5E)/LexPR,LexOP(pME)/asc-mKate2(p3E)*. Expression was induced upon addition of 10 μΜ Mifepristone (RU486, Sigma-Aldrich).

#### Site-directed mutagenesis

For site-directed mutagenesis of the *HSE:asc-mKate2* the QuikChange II XL Site-Directed Mutagenesis Kit (Agilent Technologies) was used according to manufacturer’s instructions. To make *HSE:asc-mKate2 asc* ATG morpholino-resistant a total of 6 bp changes were made with two rounds of site-directed mutagenesis, the first introduced the G6A, A9G, T12A mutations with one complementary primer pair (GCTTGAATTCACCATGGCAGAGTCATTCAAGGAGCAGCTGCAG) and the second introduced the G18A, G21A, G24A mutations (CTCAAAAGCCTCCTGCAGTTGTTCTTTGAATGACTCTGCCATGGTG). Specific primer pairs were used to mutate each phosphorylation site: T38A (GGAGGCAGGAACCGCGCGTCGCAAAGTCTGCAATCGAAAAGCTG), Y152F (CATCACAAATGAGGATTTCTGTACCATTCGTAATAAG), T160A (CCATTCGTAATAAGGAGGCTCCTCAAAAGAAGATG), T170A (GAGAGAGTTATTAGCAGGCCCAATCACATG).

#### sgRNA and mRNA synthesis

To synthesize the templates for sgRNAs targeting *caspa*, the two-oligo PCR method (10) was used. For sgRNAs targeting *asc*, sgRNA-containing plasmids were cloned using oligo annealing (11). All sgRNAs were transcribed using the MEGAshortscript T7 Transcription Kit (Ambion). To synthesize mRNA, linearized pCS2 + DNA vector containing the gene of interest was used as template and transcribed with the mMessage mMachine SP6 Transcription Kit (Ambion). RNA from *in vitro* transcriptions was purified with the RNA Clean & Concentrator-5 (Zymo Research). mRNAs were injected into one-cell stage embryos.

#### RNA extraction, cDNA synthesis and RT-PCR

Total RNA was extracted from larvae using TriFast (Peqlab) according to manufacturer’s instructions. To prevent contamination from gDNA, samples were treated with RQ1 RNase-Free DNase (Promega) and then repurified using TriFast. To generate first strand cDNA from total extracted RNA was generated using the Superscript III Reverse Transcriptase enzyme (Invitrogen). The obtained cDNA was directly used for reverse transcription PCR using Phusion High-Fidelity DNA Polymerase (Thermo Fisher Scientific). The following primers were used: *asc* (Fwd: AGTAGCAGATGATCTATTGAGG, Rev: AGAGCATCATACAAGACTTCTTTCC), *caspa* (Fwd: CAGTCAGCGCCCTGAGCTAAACATG, Rev: TCAACTGAGCTGGATCCTTCGG), *efla* (Fwd: CTTCTCAGGCTGACTGTGC, Rev: CCGCTAGCATTACCCTCC).

#### Whole-mount in situ hybridization, plastic embedding and sectioning

*In situ* hybridization was performed essentially as described previously (12). Antisense and sense probes for *asc* and *caspa* CDS were transcribed *in vitro* from linearized pCS2 + DNA vector containing the entire CDS of each gene by use of the DIG RNA Labeling Kit (Roche) and purified SigmaSpin Post-Reaction Clean-Up Columns (Sigma-Aldrich). BM Purple AP substrate (Roche) was used for staining. Whole-mount *in situ* samples were sectioned using the Leica Historesin embedding kit (Leica Microsystems) according to manufacturer’s instructions. Sectioning was carried out manually using Leica RM2235 Manual Rotary Microtome (Leica Microsystems).

#### ASC polyclonal antibody production

ASC polyclonal antibody was generated from the full-length recombinant ASC purified from a bacterial expression system. Antigen production and antibody purification were carried out by the Protein Expression and Purification Core Facility at EMBL. The rabbit immunization procedure and all animal handling were performed by the Polyclonal Antibody service at the EMBL Laboratory Animal Resources. Antibody specificity was confirmed by using preimmunization serum as a negative control and in the immunostaining pattern in *asc* morphant embryos.

#### Immunostaining

Two variants of immunostainings were used, depending on the tissue of interest. Immunostainings of myeloid cells were carried out as previously described (13). To visualize keratinocyte stainings, a less abrasive protocol lacking methanol dehydration, proteinase K treatment and postfixation steps, was used for epidermis preservation. The following primary antibodies were used antiASC (1:10^3^ dilution), antiGFP (Santa Cruz, 1:10^4^ dilution) or antiLamin B2 (1:200 dilution, Thermo Fisher Scientific). Secondary antibodies (Invitrogen) were coupled to Alexa-488, -568 and -647 (1:500, 1:500, 1:300 dilutions, respectively).

#### Protein extraction and western blotting

To obtain whole-embryo protein lysate, embryos were sonicated in fresh buffer (10 mM HEPES pH 7.5, 100 mM KCl, 2 mM MgCl_2_, 0.1 mM CaCl_2_, 5 mM EGTA pH 8.0, 1 mM NaF, 1 mM Na_3_VO_4_, 0.5% Triton, Protease inhibitor cocktail tablets [1 tablet/10 ml, Roche]). Lysate was cleared by centrifugation and supernatant was collected and stored after addition of 5xSDSSample Buffer (10% SDS, 20% glycerol, 0.2 M Tris-HCl pH 6.8, 0.05% Bromophenol Blue and 10% β-mercaptoethanol added right before use). Prepared protein samples were separated by SDS-PAGE using the Mini-PROTEAN Vertical Electrophoresis Cell system (Bio-Rad), transferred to a polyvinylidene difluoride (PVDF) membrane (Immobilion-P) in a semi-dry transfer cell (Bio-Rad) and probed using antiASC (1:10^3^ or 1:10^4^ dilution) or antiGFP (Santa Cruz, 1:10^4^ dilution) and developed with corresponding HRP-coupled secondary antibodies (Jackson ImmunoResearch). Detection was carried out using Luminata Crescendo Western HRP Substrate (Millipore).

#### Generation of caspa mutant

*sgRNA design:* Small guide RNAs (sgRNAs) targeting the first exon of zebrafish gene *caspa* (ENSDARG00000008165) were designed using the tool at http://crispr.mit.edu (14) and selected as reported (10). *sgRNA in vivo validation:* To test whether sgRNAs were targeting the region of interest *in vivo*, sgRNAs were injected in varying concentrations (120-275 ng/μl) together with 1 μl of in-house (Protein Expression and Purification facility, EMBL Heidelberg) synthesized Cas9 protein (4 mg/ml) complemented with ca. 150 mM KCl into fertilized eggs at the one-cell stage of the zebrafish TLF strain. Successful knockdown was verified by sequencing of an 800 bp PCR product from the targeted region of *caspa* (Fwd: TGGGTTAACTAGGCAAGTCAGGG, Rev: AGGGTGTATCAGGACTTGGGCCC or Rev: CCACACATGGGAGGTGTGAA). *Screening:* Embryos injected with the most efficient sgRNA (GGACGCTTTAAGTAATATTGGGG) were raised to adulthood to obtain the F0. At 6 wpf, F0 fish were genotyped by fin clipping. F0 fish showing successful targeting were incrossed and the F1 generation was raised to adulthood. Through genotyping of the F1 adults, two KO alleles were found: the *caspa*^K**^ allele carrying a 5’-AAATAATAA -3’ insertion at the expected Cas9 cleavage site resulting in two STOP codons and the *caspa*^Δ800^, carrying a deletion of ca. 800 bp including most of the first exon and part of the first intron that resulted in a nonsense mutation. Heterozygous F1 fish carrying both alleles were incrossed to obtain homozygous mutants with either the *caspa*^K**^ or the *caspa*^Δ800^ deletion allele.

#### CLEM

For CLEM analysis, the embryos were high-pressure frozen (HPM010 AbraFluid), using 20% dextran or 20% ficoll as cryoprotectant. The embryos were pierced with a needle in a cryo-microtome chamber (Leica EM FC6) at -160°C to facilitate freeze substitution (15). Embryos were then freeze-substituted (EM-AFS2, Leica Microsystems) with 0.1% Uranyl Acetate (UA) in acetone at -90°C for 48 hours. The temperature was then raised to -45°C at 3.5°C/h and samples were further incubated for 5 hours. After rinsing in acetone, the samples were infiltrated in Lowicryl HM20 resin, while raising the temperature to -25°C and left to polymerize under UV light for 48 hours at -25°C and for further 9 hours while the temperature was gradually raised to 20°C (5°C/h). Thick sections (300 nm) were cut from the polymerized resin block and picked up on carbon coated mesh grids. The imaging of sections by fluorescence microscopy (FM) was carried out as previously described (16, 17)using a widefield fluorescence microscope (Nikon TiE). Images were collected with mCherry-specific settings as well as transmitted light. TEM tomography was acquired with a FEI Tecnai F30 electron microscope. Dual-axis tomograms were obtained using SerialEM (18) and reconstructed in eTomo, part of the IMOD software package (19)(Boulder Laboratory, University of Colorado). Correlation between light and electron micrographs was carried out with the plugin ec-CLEM (http://icy.bioimageanalysis.org/plugin/ec-CLEM) of the software platform Icy (20). Features visible in both the light and electron microscopy images were manually assigned by clicking. The coordinates of pairs in the two imaging modalities were used to calculate a linear transformation, which allowed to map the coordinates of the fluorescent spot of interest (red channel) and to overlay it on the electron micrograph. The tomograms were threshold segmented with the Microscopy Image Browser platform (21), the resulting model was loaded into the digital space of Amira for visualisation (FEI Company, Hillsboro, Oregon).

#### Software

The software Geneious Version 6.1.7r was used for cloning strategy design, sequencing data analysis and sequence alignments. The kinase-specific prediction of phosphorylation sites in zebrafish ASC was carried out using the online software GPS 2.1.1 (22), using previously described parameters (23). The software Prism Version 6.03 (GraphPad) was used for all statistical analyses and graphs. Raw images were processed using ImageJ/Fiji (NIH) and Imaris x64 7.6.4 (Bitplane, AG).

---

**Table S1.** Results of JNK and Syk kinase-specific phosphorylation site prediction in zebrafish ASC by the online software GPS 2.1.1 (22).

| Position | Code | Kinase | Peptide | Score |
| --- | --- | --- | --- | --- |
| 38 | T | CMGC/MAPK/JNK/MAPK9 | RRQEPRVTKSAIEKL | 2.211 |
| 40 | S | CMGC/MAPK/JNK | QEPRVTKSAIEKLKD | 1.354 |
| 40 | S | CMGC/MAPK/JNK/MAPK9 | QEPRVTKSAIEKLKD | 3 |
| 40 | S | CMGC/MAPK/JNK/MAPK10 | QEPRVTKSAIEKLKD | 4 |
| 93 | S | CMGC/MAPK/JNK | RNTGQSESKGAPSDE | 1.333 |
| 93 | S | CMGC/MAPK/JNK/MAPK10 | RNTGQSESKGAPSDE | 3.857 |
| 152 | Y | TK/Syk | KVITNEDYCTIRNKE | 1.892 |
| 152 | Y | TK/Syk/Syk | KVITNEDYCTIRNKE | 2.627 |
| 152 | Y | TK/Syk/ZAP70 | KVITNEDYCTIRNKE | 2.95 |
| 160 | T | CMGC/MAPK/JNK | CTIRNKETPQKKMRE | 5.104 |
| 160 | T | CMGC/MAPK/JNK/MAPK8 | CTIRNKETPQKKMRE | 14.861 |
| 160 | T | CMGC/MAPK/JNK/MAPK9 | CTIRNKETPQKKMRE | 3.053 |
| 160 | T | CMGC/MAPK/JNK/MAPK10 | CTIRNKETPQKKMRE | 6.429 |
| 170 | T | CMGC/MAPK/JNK | KKMRELLTGPITCAG | 1.521 |
| 185 | Y | TK/Syk/ZAP70 | NKGKEVLYDALRESN | 2 |

**Fig. S1.**
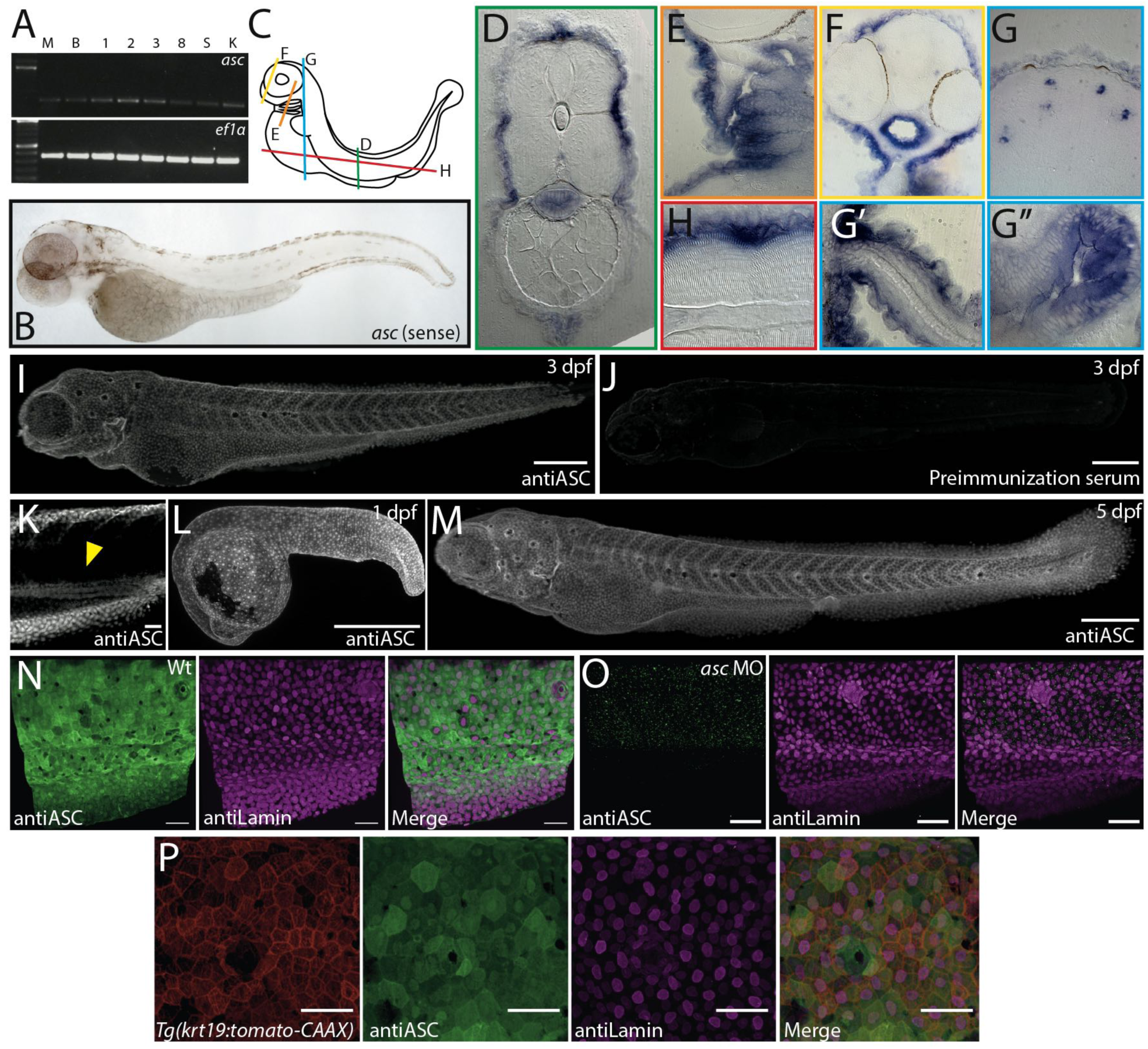
*asc* is expressed during zebrafish early development. RT-PCR of *asc* during early development in Morula (M), Blastula (B), 1, 2, 3 and 8 dpf, adult spleen (S) and adult head kidney (K). *ef1a* is used as housekeeping gene control [A]. Sense probe control for *asc wish* [B]. Diagram depicting plastic-embedded *asc* (antisense probe) *wish* sample sectioning [C] including trunk cross section [D], enlarged view of gills [E], anterior [F] and posterior [G] head region, lateral fin [G’], intestine [G”] cross sections; and of longitudinal trunk section [H]. Immunostaining in 3 dpf larvae using antiASC [I] or preimmunization serum [J]. Single plane of ASC immunostaining showing expression in intestine [yellow arrowhead, K]. Immunostainings of ASC in 1 dpf embryo [L] and 5 dpf larvae [M]. Immunostainings of ASC in 3 dpf wild type [N] and *asc* ATG-morpholino injected larvae [O]. antiLamin staining is used as positive control. Immunostaining of *Tg(krt19:tomato-CAAX)* transgenic 3 dpf larva shows ASC expression in basal and EVL keratinocytes [P]. Scale bars, 300 μm for full larvae, otherwise 50 μm.

**Fig. S2.**
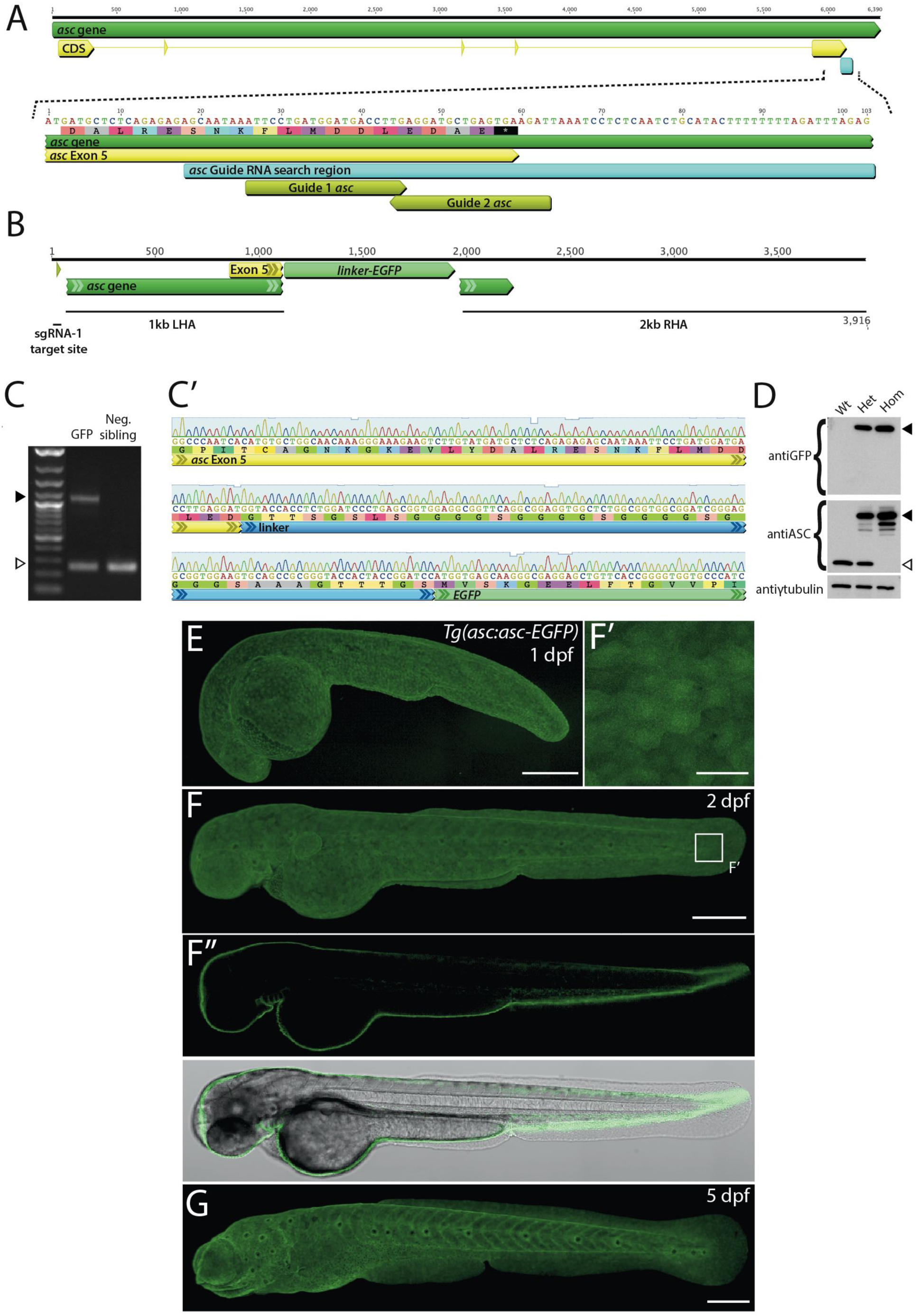
Generation, genotyping and imaging of *Tg(asc:asc-EGFP)*. Diagram of *asc* gene (green) with exons (yellow) showing sgRNA search region in final exon (teal) and Guide 1 and Guide 2 *asc* sgRNAs (lime green) [A]. Donor vector design included 1 and 2 kb left and right homology arms (LHA and RHA, respectively) flanking a linker-EGFP CDS [B]. Single F1 progeny larvae, screened based on GFP expression, were genotyped via PCR using primers flanking the Guide 2 *asc* sgRNA target site. Amplification of the wild type allele yields a 260 bp product (white arrowhead), of *asc-EGFP* allele a 1.1 kb product (black arrowhead) containing the 850 bp linker-GFP sequence [C]. Sequencing of the 1.1 kb *asc-EGFP* allele PCR product [C’]. Western blotting of full protein extracts of wild type, and heterozygous and homozygous *Tg(asc:asc-EGFP)* larvae. GFP is present only in transgenic larvae (black arrowhead). Untagged protein is absent in homozygous *Tg(asc:asc-EGFP)* larvae (white arrowhead) [D]. Live imaging of *Tg(asc:asc-gfp)* at 1 dpf [E], 2 dpf [F] and 5 dpf [G]. Magnification of epidermal cells shows ASC-GFP localization in the nucleus of epidermal cells [F’]. Optical sagittal section of 2 dpf larva with and without brightfield merge [F’’]. Scale bars, 300 gm for full larvae, otherwise 40 μm.

**Fig. S3.**
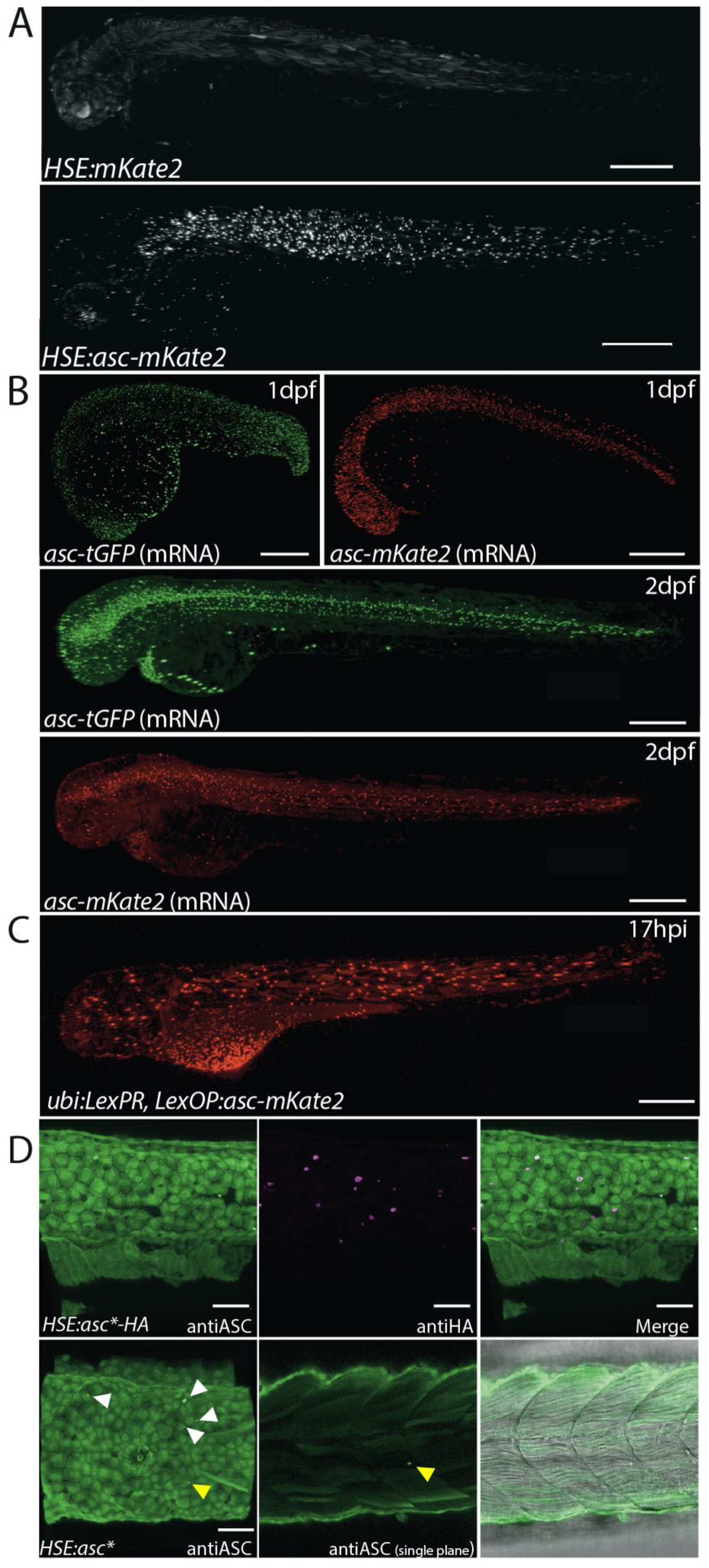
ASC misexpression *in vivo* results in speck formation. Live imaging of 3 dpf larvae transiently expressing *HSE:mKate2* or *HSE:asc-mKate2* 17 hphs [A]. Specks are only observed after heat shock in larvae expressing ASC-mKate2. Wild type *asc-mKate2* or *asc-tGFP* mRNA injected embryos at 1 and 2 dpf [B]. Live imaging of 3 dpf larvae transiently expressing *asc-mKate2* from a LexPR/OP construct driven by the *ubi* promoter. Specks are observed 17h after addition of Mifepristone to the media, which enables LexPR binding to the LexOP operator [C]. AntiASC immunostaining of 3 dpf larvae after transiently expressing *HSE:asc-HA* or *HSE:asc* [D]. Specks of ASC-HA are colabeled by antiHA (upper row). ASC specks (untagged) are highlighted by arrowheads (lower row). Scale bars, 300 μm for full larvae, otherwise 50 μm.

**Fig. S4.**
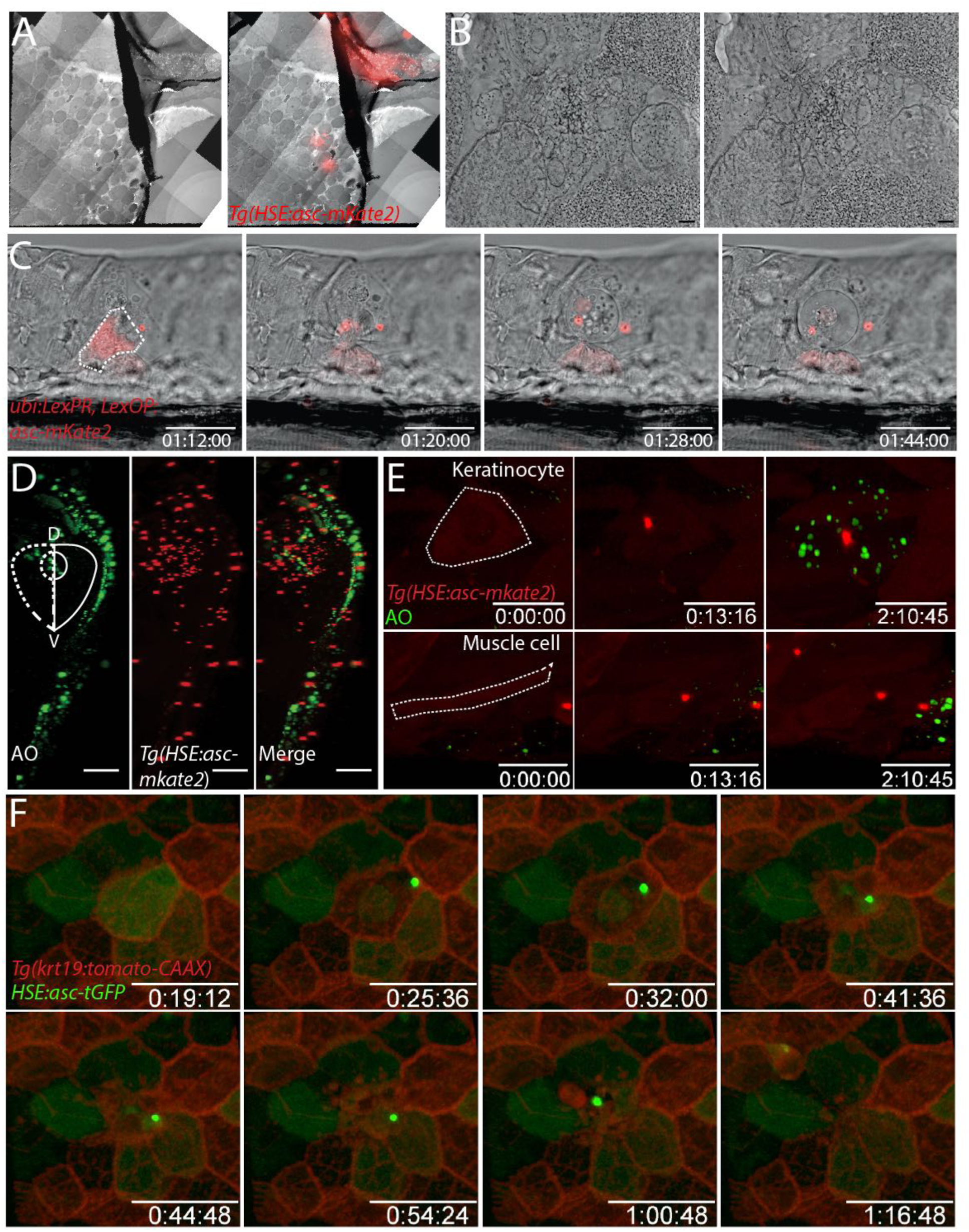
ASC speck formation leads to keratinocyte cell death. Low magnification of tissue section showing separate electron micrograph and overlay with red channel [A]. TEM tomography slices of second speck, shown in black arrowhead on fig. 4A at two different depths [B]. Scale bars, 200 nm. Live imaging of single keratinocyte transiently expressing Mifepristone-inducted ASC-mKate2 and undergoing cell death after speck formation [C]. Right side of trunk cross section of 3 dpf *Tg(HSE:asc-mKate2)* larva stained with acridine orange at 15 hphs (D, dorsal; V, ventral) [D]. Cell death mainly localizes to the epidermal layer. Time lapse imaging of single keratinocyte and muscle cell in a 3 dpf *Tg(HSE:asc-mKate2)* larva stained with acridine orange at 3 hphs [E]. Acridine orange-labeled debris accumulates only after speck formation in the keratinocyte. Time lapse imaging of 3 dpf *Tg(krt19:tomato-CAAX)* larva transiently expressing *HSE:asc-tGFP*, showing plasma membrane collapse and cell extrusion after speck formation in keratinocytes [D]. Scale bars, 30 μm.

**Fig. S5.**
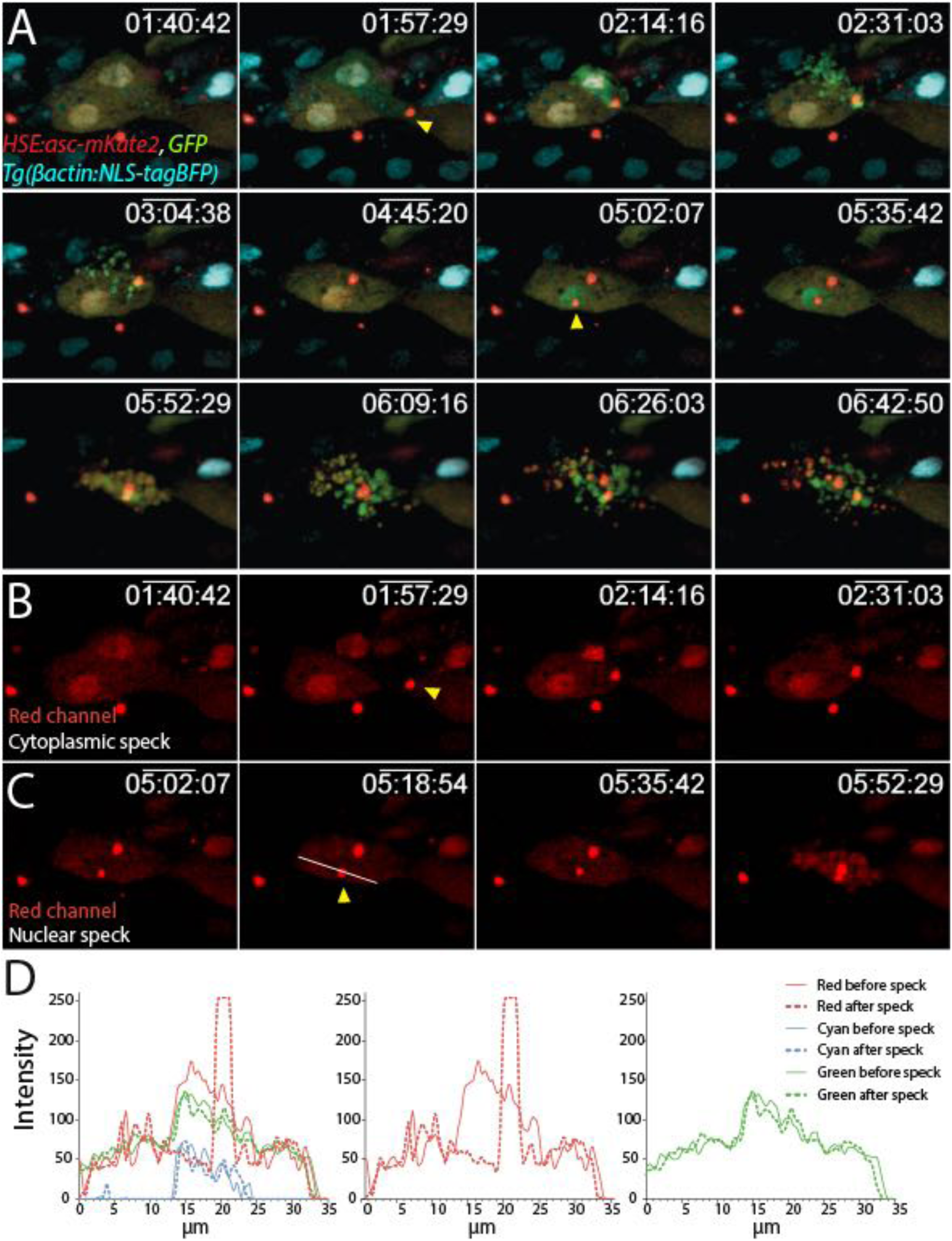
Cell death follows speck formation in the nucleus or cytoplasm of keratinocytes. Time lapse imaging of transient ASC-mKate2 and GFP expression in *Tg(βactin:NLS-tagBFP)* larvae 6 hphs [A]. Yellow arrowheads signal speck formation events in two cells; first, within the cytoplasm and second, within the nucleus. Red channel showing ASC-mKate2 depletion from cytoplasmic [B] and nuclear compartments [C] during speck formation. Intensity plot profile (white line) before and after nuclear speck formation for all channels [D]. Middle and right panels show green and red channels separately, highlighting ASC-mKate2 depletion only from nuclear pool. Scale bars, 20 μm.

**Fig. S6.**
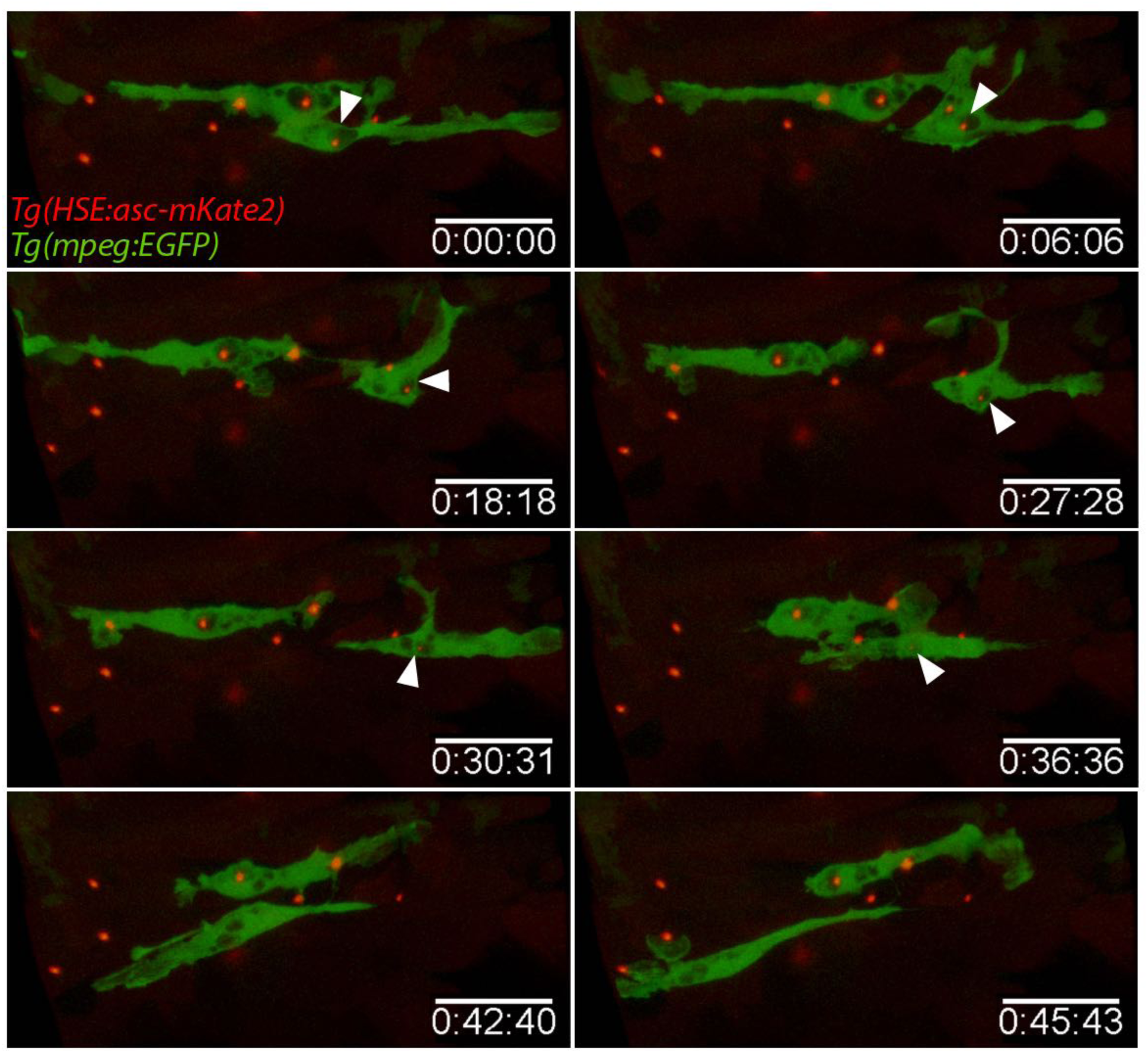
ASC specks are degraded within phagosomes. Time lapse imaging of *Tg(HSE:asc-mKate2, mpeg:EGFP)* larvae 17 hphs, showing degradation of speck within phagosome [white arrowhead]. Scale bars, 30 μm.

**Fig. S7.**
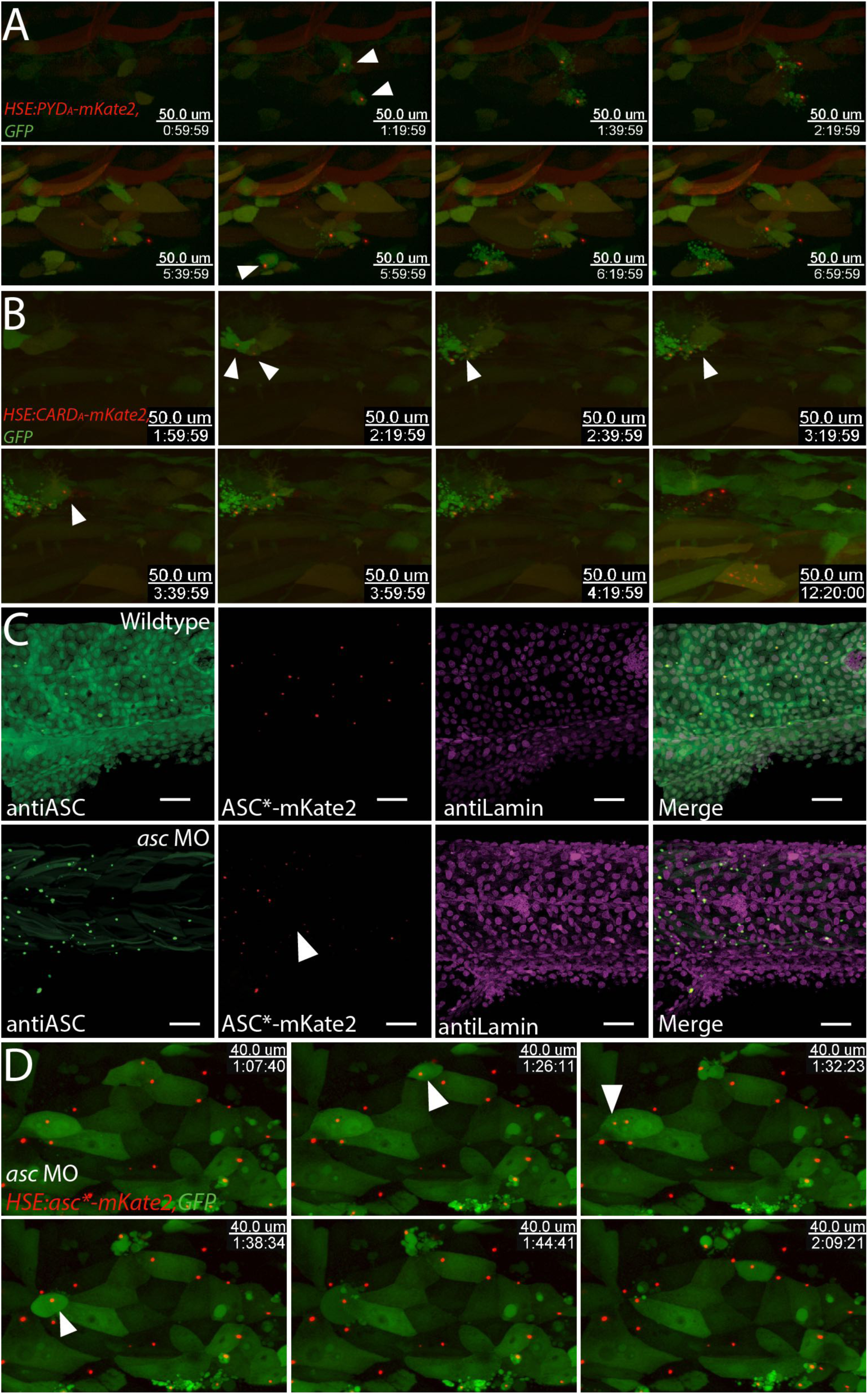
In presence of endogenous ASC, PYD_A_ or CARD_A_ overexpression leads to speck formation. Time lapse imaging of keratinocytes transiently expressing *HSE:PYD_A_-mKate2* [A] or *HSE:CARD_A_-mKate2* [B] and GFP in wild type larvae 3 hphs. Specks in keratinocytes are undistinguishable from those formed by ASC-mKate2 overexpression, and also lead to cell death (white arrowheads). Immunostaining of 3 dpf larvae expressing morpholino resistant version of *asc-mKate2* containing 6 silent mutations (*asc*-mKate2)* in wild type [C, upper row] or *asc* morpholino-injected larva [C, lower row]. Lamin is used as a positive control for the staining. Time lapse imaging of *asc* morpholino-injected *Tg(asc:asc-EGFP)* larvae transiently expressing *HSE:asc*-mKate2* with GFP [D]. Speck formation and cell death is unaffected by lack of endogenous protein. Scale bars, 20 μm.

**Fig. S8.**
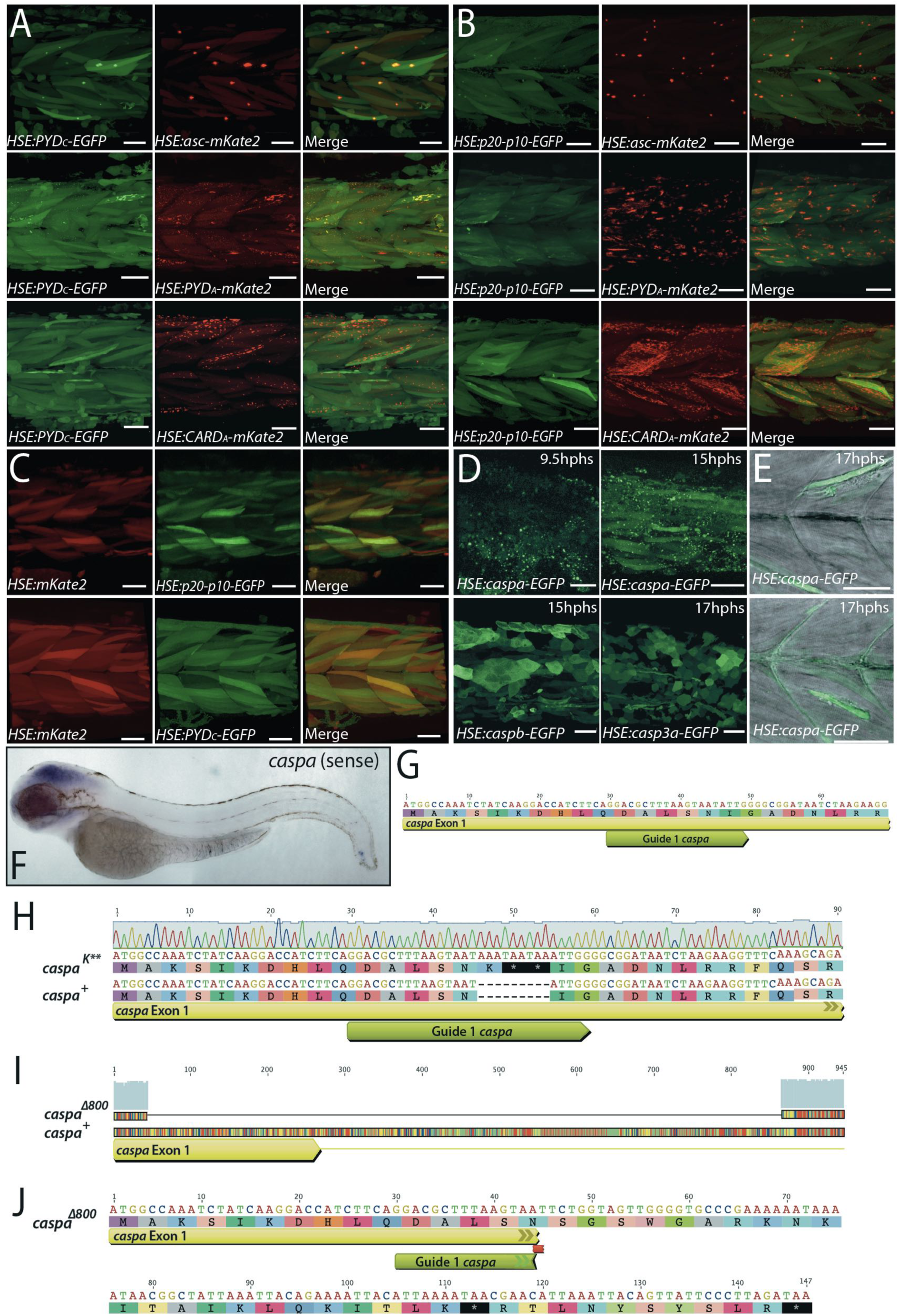
Consequences of Caspa overexpression land generation of a *caspa* mutant. Live imaging of heat-shock induced transient expression of single Caspa domains: *HSE:PYD_C_-mKate2* [A] or *HSE:p20-p10c-mKate2* [B] with full length ASC, its individual domains or mKate2 [C] around 17 hphs. Interaction only occurs when both proteins contain their respective PYD domains. Live imaging of transient expression of *HSE:caspa-EGFP, HSE:caspb-EGFP* or *HSE:casp3a-EGFP* between 9 and 17 hphs [D]. Vast amounts of epidermal cellular debris are seen only when Caspa-GFP is overexpressed. Single plane of *HSE:caspa-EGFP* transient expression 17 hphs in muscle cells showing morphological changes upon Caspa-GFP overexpression [E]. Scale bars, 50 μm. Sense probe control for *caspa wish* [F]. Generation of two *caspa* mutant alleles using CRISPR/Cas9 [G-J]. First exon of *caspa* gene (yellow) with target sites of Guide 1 *caspa* sgRNA (lime green) [G]. Sequence of *caspa^K^*\*\* allele: an insertion of 9 bp adds one lysine (K) and two STOP codons in the *caspa* reading frame [H]. Sequence of caspa^Δ800^ allele: deletion of 800 bp fragment containing 224 bp of Exon 1 and 596 bp from Intron 1 [I], causes frame shift and insertion of a STOP codon after 37 aa [J].

## Supplemental Movies

**Movie S1.** Time lapse imaging of endogenous speck formation examples in 3 dpf *Tg(asc:asc-EGFP)* larvae.

**Movie S2.** Time lapse imaging of speck formation in 3 dpf *Tg(asc:asc-EGFP)* larva induced by *HSE:NLR-mKate2* or *HSE:asc-mKate2* transient overexpression.

**Movie S3.** Time lapse imaging of speck formation in 3 dpf *Tg(HSE:asc-mKate2)* full larva in single cells.

**Movie S4.** TEM tomography stack of specks in 3 dpf *Tg(HSE:asc-mKate2)* larva 18 hphs.

**Movie S5.** Time lapse imaging of speck formation in single muscle cells of 3 dpf *Tg(HSE:asc-mKate2)* larva with brightfield, in EVL keratinocytes of 3 dpf *Tg(HSE:asc-mKate2, krt4:GFP)* larva without and with brightfield, in keratinocytes 3 dpf *Tg(HSE:asc-mKate2)* with lynGFP-labeled plasma membrane and in keratinocytes of 3 dpf *Tg(krt19:Tomato-CAAX)* transiently expressing *HSE:asc-tGFP*.

**Movie S6.** Time lapse imaging of nuclear speck formation in 3 dpf *Tg(asc:asc-EGFP)* larva expressing *HSE:NLS-asc-mKate2* and 3 dpf *Tg(βactin:NLS-tagBFP)* larvae transiently expressing *HSE:asc-mKate2* and GFP.

**Movie S7.** Time lapse imaging of *Tg(HSE:asc-mKate2, mpeg:EGFP)* larvae.

**Movie S8.** Time lapse imaging of *HSE:PYD_A_-mKate2* or *HSE:CARD_A_-mKate2* and GFP transiently expressed in 3 dpf wildtype larvae and *HSE:asc-mKate2*, *HSE:PYD_A_-mKate2* or *HSE:CARD_A_-mKate2* and GFP transiently expressed in 3 dpf *asc* morpholino-injected *Tg(asc:asc-EGFP)* larvae.

**Movie S9.** Time lapse imaging of *HSE:asc-mKate2* and GFP transiently expressed in 3 dpf wildtype or *caspa* knockout larvae.

## References

1. P. Broz, V. M. Dixit, Inflammasomes: mechanism of assembly, regulation and signalling. Nat Rev Immunol, 1–14 (2016).

2. D. Sharma, T.-D. Kanneganti, The cell biology of inflammasomes: Mechanisms of inflammasome activation and regulation. The Journal of Cell Biology. 213, 617–629 (2016).

3. A. V. Hauenstein, L. Zhang, H. Wu, The hierarchical structural architecture of inflammasomes, supramolecular inflammatory machines. Curr. Opin. Struct. Biol. 31, 75–83 (2015).

4. S. M. Man, T.-D. Kanneganti, Converging roles of caspases in inflammasome activation, cell death and innate immunity. Nat Rev Immunol. 16, 7–21 (2015).

5. J. E. Vince, J. Silke, The intersection of cell death and inflammasome activation. Cell. Mol. Life Sci. 73, 2349–2367 (2016).

6. L. Vande Walle, M. Lamkanfi, Pyroptosis. Curr Biol. 26, R568–72 (2016).

7. F. Hoss, J. F. Rodriguez-Alcazar, E. Latz, Assembly and regulation of ASC specks. Cell. Mol. Life Sci., 1–19 (2016).

8. E. de Alba, Structure and interdomain dynamics of apoptosis-associated speck-like protein containing a CARD (ASC). Journal of Biological Chemistry. 284, 32932–32941 (2009).

9. A. Lu et al., Unified Polymerization Mechanism for the Assembly of ASC-Dependent Inflammasomes. Cell. 156, 1193–1206 (2014).

10. X. Cai et al., Prion-like Polymerization Underlies Signal Transduction in Antiviral Immune Defense and Inflammasome Activation. Cell. 156, 1207–1222 (2014).

11. J. Masumoto et al., ASC, a novel 22-kDa protein, aggregates during apoptosis of human promyelocytic leukemia HL-60 cells. J Biol Chem. 274, 33835–33838 (1999).

12. T. Fernandes-Alnemri et al., The pyroptosome: a supramolecular assembly of ASC dimers mediating inflammatory cell death via caspase-1 activation. Cell Death Differ. 14, 1590–1604 (2007).

13. J. C. Kagan, V. G. Magupalli, H. Wu, SMOCs: supramolecular organizing centres that control innate immunity. Nat Rev Immunol. 14, 821–826 (2014).

14. L. Sborgi et al., Structure and assembly of the mouse ASC inflammasome by combined NMR spectroscopy and cryo-electron microscopy. Proceedings of the National Academy of Sciences. 112, 13237–13242 (2015).

15. M. S. Dick, L. Sborgi, S. Rühl, S. Hiller, P. Broz, ASC filament formation serves as a signal amplification mechanism for inflammasomes. Nat Commun. 7, 11929 (2016).

16. F. I. Schmidt et al., A single domain antibody fragment that recognizes the adaptor ASC defines the role of ASC domains in inflammasome assembly. J Exp Med. 213, 771–790 (2016).

17. J. Cheng et al., Kinetic properties of ASC protein aggregation in epithelial cells. J. Cell. Physiol. 222, 738–747 (2010).

18. D. P. Sester et al., A novel flow cytometric method to assess inflammasome formation. The Journal of Immunology. 194, 455–462 (2015).

19. A. Stutz, G. L. Horvath, B. G. Monks, E. Latz, ASC speck formation as a readout for inflammasome activation. Methods Mol Biol. 1040, 91–101 (2013).

20. M. Beilharz, D. D. Nardo’s, E. Latz, B. S. Franklin, Measuring NLR Oligomerization II: Detection of ASC Speck Formation by Confocal Microscopy and Immunofluorescence. Methods Mol Biol. 1417, 145–158 (2016).

21. T.-C. Tzeng et al., A Fluorescent Reporter Mouse for Inflammasome Assembly Demonstrates an Important Role for Cell-Bound and Free ASC Specks during In Vivo Infection. CellReports. 16, 571–582 (2016).

22. A. S. Yazdi, S. K. Drexler, J. Tschopp, The Role of the Inflammasome in Nonmyeloid Cells. J. Clin. Immunol. 30, 623–627 (2010).

23. P. M. Peeters, E. F. Wouters, N. L. Reynaert, Immune Homeostasis in Epithelial Cells: Evidence and Role of Inflammasome Signaling Reviewed. Journal of Immunology Research. 2015, 1–15 (2015).

24. P. Santana et al., Is the inflammasome relevant for epithelial cell function? Microbes Infect., 1–9 (2015).

25. L. Feldmeyer, S. Werner, L. E. French, H.-D. Beer, Interleukin-1, inflammasomes and the skin. European Journal of Cell Biology. 89, 638–644 (2010).

26. S. K. Drexler et al., Tissue-specific opposing functions of the inflammasome adaptor ASC in the regulation of epithelial skin carcinogenesis. Proceedings of the National Academy of Sciences. 109, 18384–18389 (2012).

27. S. A. Renshaw, N. S. Trede, A model 450 million years in the making: zebrafish and vertebrate immunity. Dis Model Mech. 5, 38–47 (2012).

28. V. Torraca, S. Masud, H. P. Spaink, A. H. Meijer, Macrophage-pathogen interactions in infectious diseases: new therapeutic insights from the zebrafish host model. Dis Model Mech. 7, 785–797 (2014).

29. M. van der Vaart, A. H. Meijer, H. P. Spaink, Pathogen Recognition and Activation of the Innate Immune Response in Zebrafish. Advances in Hematology. 2012, 1–19 (2012).

30. C.-Y. Lin, C.-Y. Chiang, H.-J. Tsai, Zebrafish and Medaka: new model organisms for modern biomedical research. Journal of Biomedical Science, 1–11 (2016).

31. P. Kuri, K. Ellwanger, T. A. Kufer, M. Leptin, B. Bajoghli, A high-sensitivity, bidirectional reporter to monitor NF-κB activity in cell culture and zebrafish in real-time. J Cell Sci (2016), doi:10.1242/jcs.196485.

32. C. Stein, M. Caccamo, G. Laird, M. Leptin, Conservation and divergence of gene families encoding components of innate immune response systems in zebrafish. Genome Biol. 8, R251 (2007).

33. J. D. Hansen, L. N. Vojtech, K. J. Laing, Sensing disease and danger: A survey of vertebrate PRRs and their origins. Dev Comp Immunol. 35, 886–897 (2011).

34. K. Howe et al., Structure and evolutionary history of a large family of NLR proteins in the zebrafish. Open Biol. 6, 160009 (2016).

35. J. Masumoto et al., Caspy, a zebrafish caspase, activated by ASC oligomerization is required for pharyngeal arch development. J Biol Chem. 278, 4268–4276 (2003).

36. J. Masumoto et al., Expression of apoptosis-associated speck-like protein containing a caspase recruitment domain, a pyrin N-terminal homology domain-containing protein, in normal human tissues. J. Histochem. Cytochem. 49, 1269–1275 (2001).

37. F. Peri, C. Nüsslein-Volhard, Live imaging of neuronal degradation by microglia reveals a role for v0-ATPase a1 in phagosomal fusion in vivo. Cell. 133, 916–927 (2008).

38. P. P. Hernandez et al., Sublethal concentrations of waterborne copper induce cellular stress and cell death in zebrafish embryos and larvae. Biol. Res. 44, 7–15 (2011).

39. F. A. Olivari, P. P. Hernandez, M. L. Allende, Acute copper exposure induces oxidative stress and cell death in lateral line hair cells of zebrafish larvae. Brain Research. 1244, 1–12 (2008).

40. C. A. d'Alençon et al., A high-throughput chemically induced inflammation assay in zebrafish. BMC Biol. 8, 151 (2010).

41. B. Bajoghli, N. Aghaallaei, T. Heimbucher, T. Czerny, An artificial promoter construct for heat-inducible misexpression during fish embryogenesis. Dev Biol. 271, 416–430 (2004).

42. A. Lu, H. Wu, Structural mechanisms of inflammasome assembly. FEBS Journal. 282, 435–444 (2015).

43. H. Hara et al., Phosphorylation of the adaptor ASC acts as a molecular switch that controls the formation of speck-like aggregates and inflammasome activity. Nat Immunol. 14, 1247–1255 (2013).

44. Y.-C. Lin et al., Syk is involved in NLRP3 inflammasome-mediated caspase-1 activation through adaptor ASC phosphorylation and enhanced oligomerization. Journal of Leukocyte Biology. 97, 825–835 (2015).

45. I. Jorgensen, Y. Zhang, B. A. Krantz, E. A. Miao, Pyroptosis triggers pore-induced intracellular traps (PITs) that capture bacteria and lead to their clearance by efferocytosis. Journal of Experimental Medicine. 213, 2113–2128 (2016).

46. A. Baroja-Mazo et al., The NLRP3 inflammasome is released as a particulate danger signal that amplifies the inflammatory response. Nat Immunol. 15, 738–748 (2014).

47. B. S. Franklin et al., The adaptor ASC has extracellular and “prionoid” activities that propagate inflammation. Nat Immunol. 15, 727–737 (2014).

48. L. N. Vojtech, N. Scharping, J. C. Woodson, J. D. Hansen, Roles of Inflammatory Caspases during Processing of Zebrafish Interleukin-1β in Francisella noatunensis Infection. Infect Immun. 80, 2878–2885 (2012).

49. W. J. B. Vincent, C. M. Freisinger, P.-Y. Lam, A. Huttenlocher, J.-D. Sauer, Macrophages mediate flagellin induced inflammasome activation and host defense in zebrafish. Cellular Microbiology. 18, 591–604 (2016).

50. M. Varela et al., Cellular visualization of macrophage pyroptosis and interleukin-1β release in a viral hemorrhagic infection in zebrafish larvae. J Virol. 88, 12026–12040 (2014).

51. S. D. Tyrkalska et al., Neutrophils mediate Salmonella Typhimurium clearance through the GBP4 inflammasome-dependent production of prostaglandins. Nat Commun. 7, 12077 (2016).

52. S. Rakers et al., “Fish matters”: the relevance of fish skin biology to investigative dermatology. Exp. Dermatol. 19, 313–324 (2010).

53. M. Pasparakis, I. Haase, F. O. Nestle, Mechanisms regulating skin immunity and inflammation. Nat Rev Immunol. 14, 289–301 (2014).

54. L. Feldmeyer et al., The Inflammasome Mediates UVB-Induced Activation and Secretion of interleukin-1β by Keratinocytes. Current Biology. 17, 1140–1145 (2007).

55. H. Watanabe et al., Activation of the IL-1β-Processing Inflammasome Is Involved in Contact Hypersensitivity. J Investig Dermatol. 127, 1956–1963 (2007).

56. M. Reinholz et al., HPV16 activates the AIM2 inflammasome in keratinocytes. Arch Dermatol Res. 305, 723–732 (2013).

57. X. Dai et al., Mite allergen is a danger signal for the skin via activation of inflammasome in keratinocytes. J. Allergy Clin. Immunol. 127, 806–14.e1–4 (2011).

58. E. M. Weinheimer-Haus, R. E. Mirza, T. J. Koh, Nod-like receptor protein-3 inflammasome plays an important role during early stages of wound healing. PLoS ONE. 10, e0119106 (2015).

59. X. Cai, H. Xu, Z. J. Chen, Prion-Like Polymerization in Immunity and Inflammation. Cold Spring Harb Perspect Biol, a023580 (2016).

60. B. Balci-Peynircioglu et al., Expression of ASC in Renal Tissues of Familial Mediterranean Fever Patients with Amyloidosis: Postulating a Role for ASC in AA Type Amyloid Deposition. Experimental Biology and Medicine. 233, 1324–1333 (2008).

61. P. Sagoo et al., In vivo imaging of inflammasome activation reveals a subcapsular macrophage burst response that mobilizes innate and adaptive immunity. Nat Med. 22, 64–71 (2016).

62. M. Stemmer, T. Thumberger, M. Del Sol Keyer, J. Wittbrodt, J. L. Mateo, CCTop: An Intuitive, Flexible and Reliable CRISPR/Cas9 Target Prediction Tool. PLoS ONE. 10, e0124633 (2015).

63. W. Y. Hwang et al., Efficient genome editing in zebrafish using a CRISPR-Cas system. Nat Biotechnol. 31, 227–229 (2013).

64. S. Kirchmaier, K. Lust, J. Wittbrodt, Golden GATEway cloning--a combinatorial approach to generate fusion and recombination constructs. PLoS ONE. 8, e76117 (2013).

65. Y. Hisano et al., Precise in-frame integration of exogenous DNA mediated by CRISPR/Cas9 system in zebrafish. Sci Rep. 5, 8841 (2015).

## Supplemental References

1. M. Westerfield, The Zebrafish Book (2007).

2. F. Ellett, L. Pase, J. W. Hayman, A. Andrianopoulos, G. J. Lieschke, mpegl promoter transgenes direct macrophage-lineage expression in zebrafish. Blood. 117, e49–56 (2011).

3. D. Sieger, C. Moritz, T. Ziegenhals, S. Prykhozhij, F. Peri, Long-range Ca2+ waves transmit brain-damage signals to microglia. Dev Cell. 22, 1138–1148 (2012).

4. C. Hall, M. V. Flores, T. Storm, K. Crosier, P. Crosier, The zebrafish lysozyme C promoter drives myeloid-specific expression in transgenic fish. BMC Dev Biol. 7, 42 (2007).

5. B. Fischer et al., p53 and TAp63 promote keratinocyte proliferation and differentiation in breeding tubercles of the zebrafish. PLoS Genet. 10, e1004048 (2014).

6. F. Peri, C. Nüsslein-Volhard, Live imaging of neuronal degradation by microglia reveals a role for v0-ATPase al in phagosomal fusion in vivo. Cell. 133, 916–927 (2008).

7. B. Bajoghli, N. Aghaallaei, T. Heimbucher, T. Czerny, An artificial promoter construct for heat-inducible misexpression during fish embryogenesis. Dev Biol. 271, 416–430 (2004).

8. K. M. Kwan et al., The Tol2kit: a multisite gateway-based construction kit for Tol2 transposon transgenesis constructs. Dev. Dyn. 236, 3088–3099 (2007).

9. A. Emelyanov, S. Parinov, Mifepristone-inducible LexPR system to drive and control gene expression in transgenic zebrafish. Dev Biol. 320, 113–121 (2008).

10. A. N. Shah, C. F. Davey, A. C. Whitebirch, A. C. Miller, C. B. Moens, Rapid reverse genetic screening using CRISPR in zebrafish. Nat Meth (2015), doi:10.1038/nmeth.3360.

11. M. Stemmer, T. Thumberger, M. Del Sol Keyer, J. Wittbrodt, J. L. Mateo, CCTop: An Intuitive, Flexible and Reliable CRISPR/Cas9 Target Prediction Tool. PLoS ONE. 10, e0124633 (2015).

12. C. Thisse, B. Thisse, High-resolution in situ hybridization to whole-mount zebrafish embryos. Nat Protoc. 3, 59–69 (2008).

13. M. Varela et al., Cellular visualization of macrophage pyroptosis and interleukin-1β release in a viral hemorrhagic infection in zebrafish larvae. J Virol. 88, 12026–12040 (2014).

14. P. D. Hsu et al., DNA targeting specificity of RNA-guided Cas9 nucleases. Nat Biotechnol. 31, 827–832 (2013).

15. M. Eltsov et al., Quantitative analysis of cytoskeletal reorganization during epithelial tissue sealing by large-volume electron tomography. Nat Cell Biol. 17, 605–614 (2015).

16. O. Avinoam, M. Schorb, C. J. Beese, J. A. G. Briggs, M. Kaksonen, ENDOCYTOSIS. Endocytic sites mature by continuous bending and remodeling of the clathrin coat. Science. 348, 1369–1372 (2015).

17. W. Kukulski et al., Correlated fluorescence and 3D electron microscopy with high sensitivity and spatial precision. The Journal of Cell Biology. 192, 111–119 (2011).

18. D. N. Mastronarde, Automated electron microscope tomography using robust prediction of specimen movements. J. Struct. Biol. 152, 36–51 (2005).

19. J. R. Kremer, D. N. Mastronarde, J. R. McIntosh, Computer visualization of three-dimensional image data using IMOD. J. Struct. Biol. 116, 71–76 (1996).

20. F. de Chaumont et al., Icy: an open bioimage informatics platform for extended reproducible research. Nat Meth. 9, 690–696 (2012).

21. I. Belevich, M. Joensuu, D. Kumar, H. Vihinen, E. Jokitalo, Microscopy Image Browser: A Platform for Segmentation and Analysis of Multidimensional Datasets. Plos Biol. 14, e1002340 (2016).

22. Y. Xue et al., GPS 2.1: enhanced prediction of kinase-specific phosphorylation sites with an algorithm of motif length selection. Protein Eng. Des. Sel. 24, 255–260 (2011).

23. H. Hara et al., Phosphorylation of the adaptor ASC acts as a molecular switch that controls the formation of speck-like aggregates and inflammasome activity. Nat Immunol. 14, 1247–1255 (2013).

